# Mapping of safe and early chemo-attenuated live *Plasmodium falciparum* immunization identifies immune signature of vaccine efficacy

**DOI:** 10.1101/2020.09.14.296152

**Authors:** Steffen Borrmann, Zita Sulyok, Katja Müller, Rolf Fendel, Carlos Lamsfus Calle, Mihaly Sulyok, Johannes Friesen, Albert Lalremruata, Thaisa Lucas Sandri, The Trong Nguyen, Annette Knoblich, Stephanie Sefried, Javier Ibáñez, Freia-Raphaella Lorenz, Henri Lynn Heimann, David M. Weller, Regina Steuder, Selorme Adukpo, Patricia Granados Bayon, Zsófia Molnár, Meral Esen, Wolfram Metzger, Eric. R. James, Adam Ruben, Yonas Abebe, Sumana Chakravarty, Anita Manoj, KC Natasha, Tooba Murshedkar, Julius C.R. Hafalla, Tamirat Gebru Woldearegai, Fiona O’Rourke, Jana Held, Pete Billingsley, B. Kim Lee Sim, Thomas L. Richie, Stephen L. Hoffman, Peter G. Kremsner, Kai Matuschewski, Benjamin Mordmüller

**Affiliations:** Institut für Tropenmedizin, Eberhard Karls University Tübingen, Germany; German Center for Infection Research (DZIF), partner site Tübingen, Germany; Centre de Recherches Médicales de Lambaréné, Gabon; Dept. of Molecular Parasitology, Institute of Biology, Humboldt University, Berlin, Germany; Sanaria Inc., Rockville, MD, USA; Protein Potential LLC, Rockville, MD, USA; Department of Immunology and Infection, Faculty of Infectious and Tropical Diseases, London School of Hygiene and Tropical Medicine, London, United Kingdom

## Abstract

Potent protection against malaria can be induced by attenuated live-immunization with *Plasmodium falciparum* (Pf) sporozoites (SPZ). However, a better understanding of the critical processes involved in the establishment of protective immunity is needed. We explored the safety and vaccine efficacy of early chemo-attenuation of PfSPZ under atovaquone-proguanil (AP). AP caused early arrest of *P. berghei* liver stages. Despite the absence of replication, robust protection in mice correlated with parasite-specific effector-memory CD8^+^ T-cell responses. In a phase I clinical trial a single dose of AP prevented Pf infections in the liver of adult, human subjects who received three doses of 5.12x10^4^ or 1.5x10^5^ PfSPZ by direct venous inoculation combined with oral AP. However, only 2 of 8 (25%) and 2 of 10 (20%), respectively, were protected against controlled human malaria infection (CHMI) 10 weeks after the last vaccine dose, despite levels of IgG antibodies to the Pf circumsporozoite protein (PfCSP) comparable to those achieved in fully protected volunteers after immunization with 5.12x10^4^ PfSPZ with chloroquine chemoprophylaxis active only against subsequent blood stages. We identify lower IgG recognition of the secreted liver stage-specific antigens LISP2 and LSA1 and the multi-stage antigen MSP5 as immune signatures of inferior vaccine efficacy compared to PfSPZ with chloroquine chemoprophylaxis. In conclusion, we show that immune signatures of liver stage antigens, but neither an established rodent malaria model nor concentrations of antibodies against the major surface protein of sporozoites, permit prediction of vaccine efficacy. Thus, this study provides a clear rationale for the development of live sporozoite vaccination protocols that boost exposure to Pf liver stage antigens.

**Significance Statement:** Our research demonstrates that attenuation of liver infection of high doses of *Plasmodium falciparum* sporozoites by concomitant single-dose administration of atovaquone-proguanil is safe in humans. However, vaccine efficacy was modest when compared to an identical protocol using chloroquine that acts only on the subsequent blood infection. Immune signatures of secreted *P. falciparum* liver stage antigens, but neither an established rodent malaria model nor concentrations of sporozoite antibodies, permit prediction of vaccine efficacy.

## Introduction

An unprecedented global effort to fight malaria in all subtropical and tropical regions of the world has considerably reduced the disease burden. Progress, however, has stalled at an estimated 200 million cases and 450,000 deaths globally per year (1). Further advances towards elimination are likely to be facilitated only by an efficient vaccine that complements the existing conventional control measures such as vector control, long-lasting impregnated bednets and access to rapid diagnosis and chemotherapy (2).

Malaria is caused by single-cell eukaryotes of the genus *Plasmodium*. Morbidity and mortality result from a rapid, asexual expansion phase in the blood. In the case of *Plasmodium falciparum* (Pf), which accounts for nearly all malaria-related deaths, blood stage parasite biomass can be substantial and can reach up to 10^12^ infected red blood cells. Asexual blood stage infection, however, is preceded by a small mosquito-transmitted inoculum of ∼10-400 Pf sporozoites (SPZ), which specifically invade hepatocytes and replicate therein (3). Because this 1-week liver phase of infection (i) is clinically silent and (ii) represents a life-cycle bottleneck, it provides an early target for a malaria vaccine that would prevent blood stage infection with malaria parasites and thereby prevent all disease and transmission stages (4–9).

Currently there is no vaccine against malaria parasites or any other eukaryotic human pathogen, which has received marketing authorization (licensure) by the European Medicines Agency (EMA) or the U.S. Food and Drug Administration (FDA). However, a partially effective sub-unit vaccine against malaria (RTS,S/AS01E) (10) has received a favorable scientific opinion by the EMA regarding its quality, safety and short-term efficacy (11).

Superior vaccine efficacy (VE) against Pf malaria as compared to results reported for subunit vaccines has been demonstrated by intravenous inoculation of radiation-attenuated, aseptic, purified, cryopreserved PfSPZ, Sanaria^®^ PfSPZ Vaccine (6, 12–15), and intravenous inoculation of infectious, aseptic, purified, cryopreserved PfSPZ (Sanaria^®^ PfSPZ Challenge) with concomitant chemoprophylaxis with chloroquine, Sanaria^®^ PfSPZ-CVac (CQ) (7, 16). Importantly, initial results indicate higher, per-sporozoite VE of PfSPZ-based vaccine protocols that rely on late chemo-attenuation such as those using the chemoprophylactic drug chloroquine (CQ) which is active only against blood stage parasites (7, 17). Based on this, it has been postulated that the parasite expansion in the liver substantially boosts vaccine potency (7, 17), compared to radiation-attenuated parasites that undergo developmental arrest soon after hepatocyte invasion. This strategy however requires that adequate drug concentrations are maintained in the blood beyond the liver stage period in order to kill the parasites when they emerge from the liver, e.g., by weekly administration of the antimalarial drug chloroquine (CQ). It also entails the safety risk of exposing the vaccinee to transient parasitemia and mild symptoms of malaria 7-9 days after immunization. To increase the safety margin of live-immunizations it would be preferable if the parasites never emerged from the liver without compromising VE. Moreover, chemoprophylaxis that acts against liver stages may substantially strengthen a vaccination regimen with concomitant administration of PfSPZ and chemoprophylaxis and thus, would be a significant step towards a simplified, pragmatic PfSPZ immunization protocol.

Here, we present the results of pre-clinical experiments and a clinical trial exploring the VE and mechanism of protection of PfSPZ co-administered with the chemoprophylactic drug combination atovaquone-proguanil (AP). This licensed drug combination was used because in contrast to CQ, which kills Pf parasites only in infected red blood cells, AP also kills Pf parasites in the liver before they can emerge into the bloodstream (18–20).

## Results

### Exposure to atovaquone-proguanil leads to early *Plasmodium berghei* liver stage arrest

To initiate our pre-clinical analysis, we determined the *in vitro* effect of a single dose of atovaquone (A) alone or AP when added to cultured hepatoma cells together with *Plasmodium berghei* (Pb) sporozoites (Fig. 1A). Quantification of liver stage volume, morphology, and number revealed small, developmentally arrested liver stages (Fig. 1A, Fig. S1B). Notably, A- and AP-treated parasites persisted for several days, indicative of developmental arrest rather than parasite death (Fig. S1A). Hepatocyte invasion appeared to be unaffected by drug exposure since the number of intracellular liver stages was indistinguishable from untreated controls (Fig. S1C). Causal prophylactic activity was confirmed in a murine malaria model (Fig. 1B, C, D). Administration of a single dose of 3 mg/kg atovaquone plus 1.2 mg/kg proguanil or 3 mg/kg atovaquone alone in C57BL/6 mice inoculated with 10^4^ Pb sporozoites prevented blood stage infection (Fig. 1B). Quantitative RT-PCR (qRT-PCR) analysis of steady-state levels of Pb18S rRNA and, for comparison, mouse *GAPDH* mRNA in infected livers 42h after Pb sporozoite and drug co-administration (Fig. 1C, D) revealed a >1,000-fold reduced liver stage load in drug-treated mice, in good agreement with our *in vitro* data (Fig. 1A, Fig. S1). Furthermore, when Pb sporozoites were pre-exposed to AP on ice for 2h, all C57BL/6 mice that received 10^4^ treated (*n*=5) or untreated Pb sporozoites (*n*=3) developed blood stage infection after three days, confirming earlier results of no direct effect on sporozoites (21). Thus, exposure to atovaquone or atovaquone-proguanil exerts prophylactic activity against Pb liver stage parasites, but not Pb sporozoites, allowing maximum hepatocyte invasion followed by robust early liver stage arrest.

**Fig. 1.**
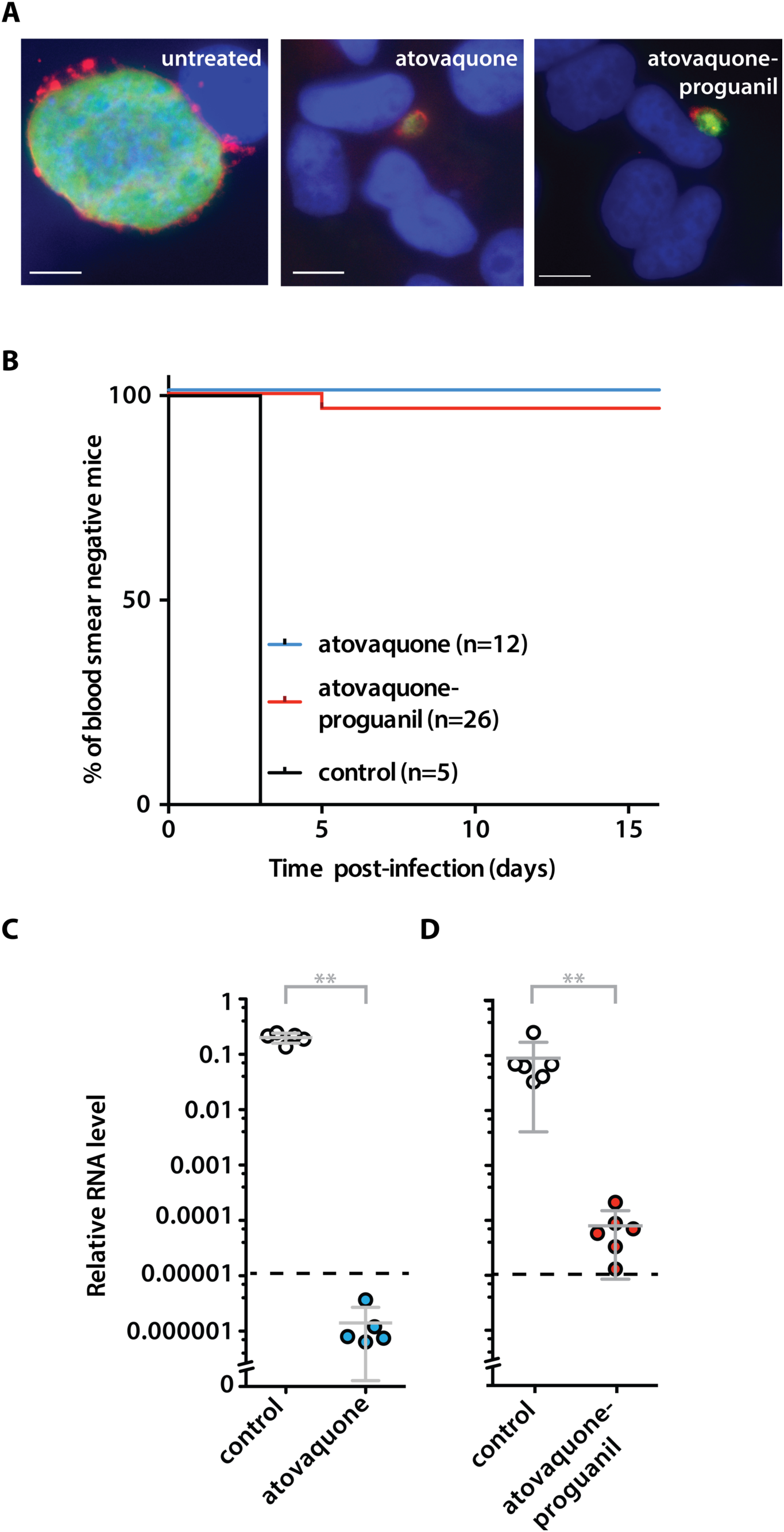
Early arrest of *Plasmodium berghei* liver stage development after co-administration of live sporozoites and atovaquone or atovaquone-proguanil. (A) Composite fluorescence micrographs of *Plasmodium berghei* (Pb) liver stages in cultured hepatoma cells. Shown are representative images of liver stages 48h after infection with sporozoites. During the first 3h cultures were exposed to atovaquone (A), atovaquone-proguanil (AP) or buffer only. Parasites were visualized by fluorescent staining of the cytoplasm (green; anti-PbHSP70 antibody), the parasitophorous vacuolar membrane (red; anti-PbUIS4 anti-serum), and nuclei (blue; Hoechst 33342). Scale bars: 10 µm. (B) Kaplan-Meier analysis of the proportion of C57BL/6 mice that remained blood film-negative after a single intravenous dose of 10^4^ *P. berghei* sporozoites without drug (black line; *n*=5) or co-administration of 3 mg/kg A (blue line) or 3/1.2 mg/kg AP (red line). Shown are cumulative data from two (A co-administration) and five (AP co-administration) independent experiments (*n*≥5 mice each). (C, D) Liver parasite load 42h after infection of C57BL/6 mice with 10^4^ sporozoites and co-administration of (C) 3 mg/kg A (blue circles) or (D) 3/1.2 mg/kg AP (red circles) or no drug (white circles). Shown are mean values (±S.D.) of relative RNA levels of Pb18S rRNA normalized to mouse *GAPDH* (*n*≥5). **, *p*< 0.01 (Mann-Whitney U test)

### Pb sporozoite immunization under atovaquone (-proguanil) cover induces sterile protection and robust IFNγ-secreting CD8^+^ CD11a^+^ T-cell responses

As proof of concept, we next immunized three groups of C57BL/6 mice with 10^4^ Pb sporozoites and concomitant administration of A or AP at weekly intervals (Fig. 2). Mice were challenged 3-4 weeks after the last immunization dose by intravenous (i.v.) inoculation of 10^4^ Pb sporozoites. Three immunizations at weekly intervals with 10^4^ Pb sporozoites and atovaquone resulted in sterile protection in all (6/6) mice. Remarkably, only two immunizations with Pb sporozoites and either A or AP still elicited sterile protection in 88% (15/17) of mice and a substantial delay to blood infection in the two mice that were not protected (Fig. 2A). We independently confirmed our findings by quantification of liver stage parasites by qRT-PCR (Fig. 2B). After sporozoite challenge parasite liver loads were reduced by at least two orders of magnitude in immunized animals as compared to controls (p<0.05) (Fig. 2B). Potential residual effects of drug treatment that could have interfered with assessment of vaccine efficacy (VE) were ruled out in an independent experiment (Fig. S2).

**Fig 2.**
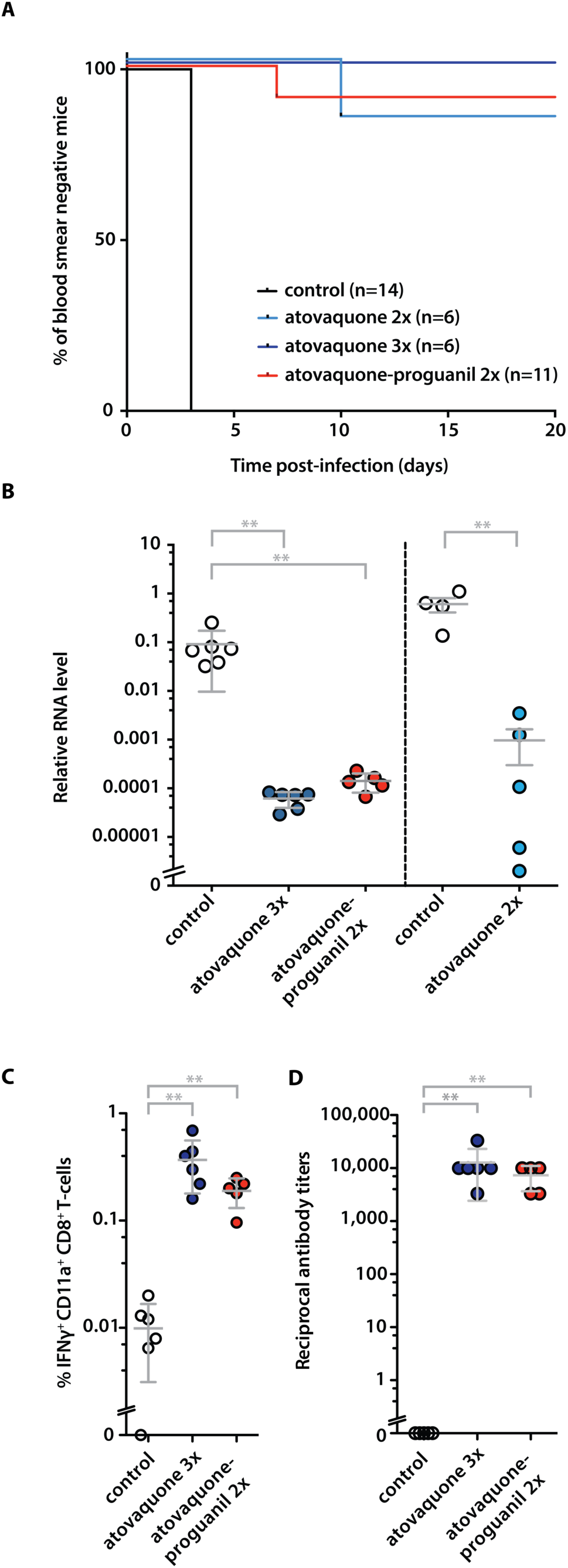
Robust protection against sporozoite challenge infections and antigen-specific immune responses after sporozoite/atovaquone (-proguanil) immunization. (A) Kaplan-Meier analysis of protection in mice immunized by co-administration of sporozoites and a single dose of atovaquone (A; 3 mg/kg i.p.) or atovaquone-proguanil (AP; 3/1.2 mg/kg i.p.). Mice were either immunized twice (AP co-administration, red line; *n*=11; A co-administration, blue line; *n*=6) or three times (A co-administration, dark blue line; *n*=6). Naïve mice served as controls (black line; *n*=14). Sporozoite challenge was done by i.v. injection of 10^4^ sporozoites three to four weeks after the last immunization. (B) Quantification of parasite liver loads after challenge infection. Mice were immunized twice (A co-administration, *n*=5, blue circles; AP co-administration; *n*= 5, red circles) and three times (A co-administration, *n*= 6, dark blue circle) as in (B). Naïve mice served as controls (*n*=6 and 4, respectively, white circles). Challenge infection was done by i.v. injection of 10^4^ sporozoites at least three weeks after the last immunization. Livers were harvested 42h later and parasite RNA quantified by RT-PCR. Relative RNA levels of Pb18S rRNA were normalized to mouse *GAPDH*. Shown are mean values (±S.D.). **: *p*< 0.01 (Mann-Whitney U test). (C) Quantification of SSP2/TRAP_130-138_ peptide-specific IFNγ-secretion by CD8^+^ CD11a^+^ T-cells from spleens of immunized or control mice (*n*≥5 each). Shown are mean values (±S.D.). **, *p*< 0.01 (Mann-Whitney U Test). (D) Quantification of anti-sporozoite antibody titers from serum of immunized or control mice (*n*≥5 each). Shown are mean values (±S.D.). **, *p*< 0.01(Mann-Whitney U Test).

We next quantified signatures of effector-memory CD8^+^ T-cell responses that correlate with protection against challenge with wild-type sporozoites (22, 23) by measuring IFNγ-secretion of CD8^+^ CD11a^+^ T-cells after stimulation with the peptides SSP2/TRAP_130-138_ and S20_318-326_, which are recognized in immunized H2-K^b^ –restricted C57BL/6 mice (24, 25) (Fig. 2C, Figs. S3 and S4). Mice were immunized with two doses of Pb sporozoites and AP or A, and lymphocytes were isolated from spleens three or four weeks after the last immunization, respectively. We detected high levels of antigen-specific IFNg secretion after both immunization protocols (Fig. 2C). Total numbers of effector memory CD8^+^ CD62L^-^ T-cells were also enhanced after immunization (Fig. S4A).

Exposure to sporozoite inoculations activates antibody-producing B cells (26–28). Accordingly, the immunization protocols also induced high (∼1:10^4^) anti-Pb sporozoite antibody titers (Fig. 2D).

In conclusion, the preclinical data demonstrate high chemoprophylactic efficacy of AP leading to reliable, early arrest of liver stage development when co-administered with Pb sporozoites. Co-administered AP with intravenous Pb sporozoites as part of an immunization protocol, resulted in parasite-specific effector-memory CD8^+^ T-cell responses and robust protection against challenge infections. Of note, vaccine efficacy was markedly superior to another early-arrest protocol with primaquine and indeed, comparable to the most potent chemo-attenuation protocols tested so far (29), prompting the design of a clinical trial.

### Clinical trial of PfSPZ-CVac (atovaquone-proguanil)

Based on the positive pre-clinical results we conducted a randomized, double-blind, placebo-controlled clinical trial of PfSPZ-CVac (AP) from September 2016 to November 2017 at the Institute for Tropical Medicine in Tübingen, Germany (Malaria controlled human infection trial E, MALACHITE; ClinicalTrials.gov Identifier: NCT02858817). The study population was selected to represent healthy, malaria-naïve volunteers aged 18–45 years from Tübingen and the surrounding area. In total, 30 volunteers (15 per dosage/group) were enrolled; 15 in Group A (PfSPZ reference dose) and 15 in Group B (PfSPZ 3-fold higher dose). They were randomly allocated to receive three injections of Sanaria® PfSPZ Challenge (aseptic, purified, cryopreserved PfSPZ of the NF54 strain, n=10 in each group) or normal saline placebo (n=5 in each group) per dosage group (30, 31).

In Group A, participants received three doses of 5.12x10^4^ PfSPZ by direct venous inoculation (DVI) at 4-week intervals and oral administration of a single dose of AP (1,000 mg/400 mg) within one hour before each PfSPZ inoculation. In Group B, participants received 1.5x10^5^ PfSPZ with the same schedule and chemoprophylactic regimen. Ten weeks after last immunization, participants in both groups underwent controlled human malaria infection (CHMI) by DVI of 3.2x10^3^ PfSPZ of PfSPZ Challenge (NF54).

Twenty-seven participants were included in the per-protocol analysis. Three withdrawals occurred, all of them in Group A before CHMI; one requested by the participant and two withdrawals by the investigators based on non-compliance or non-availability for critical study procedures. An additional participant in Group A was lost to follow-up after CHMI and was included in the per-protocol analysis. This volunteer developed parasitemia and started treatment on day 12 post-CHMI. Following successful completion of antimalarial treatment, thick blood smear (TBS) and qPCR were negative on day 21 post-CHMI. On day 22 post-CHMI the volunteer was not reachable and refused further follow-up visits. The study flow chart is shown in Fig. S5. Baseline population characteristics are detailed in Table S1.

Importantly, during immunization no breakthrough parasitemia by qPCR occurred, demonstrating robust causal prophylactic activity of a single-dose of 1,000 mg/400 mg AP, even with an inoculum of 1.5x10^5^ PfSPZ. This sporozoite dose greatly exceeds the infective dose of 3.2x10^3^ PfSPZ (31, 32) by ∼50 times and is estimated to be equivalent to the bites of ∼ 200 infected mosquitoes. Of note, safety and tolerability during immunization were similar between placebo controls and vaccinees (Table S1).

Upon challenge by CHMI, 6 out of 8 vaccinees in group A and all 4 placebo recipients developed Pf parasitemia (VE, 25%; 95% CI, -12%-50%). Due to the low efficacy and according to protocol, no heterologous repeat CHMI was performed. Subsequently, Group B underwent homologous CHMI with PfSPZ Challenge (NF54), i.e., with the vaccine strain only. In Group B, 8 of 10 vaccinees and all 5 placebo controls developed Pf parasitemia (VE, 20%; 95% CI, -9%- 41%). Moreover, time to qPCR detectable parasitemia, a marker for partial efficacy, was similar between the groups (Group A; median 7 days; IQR 7-8.5 and Group B; median 7 days; IQR 7-7.5; Kruskal-Wallis H test, p=0.27) (Fig. 3 and Fig. 4).

**Fig. 3.**
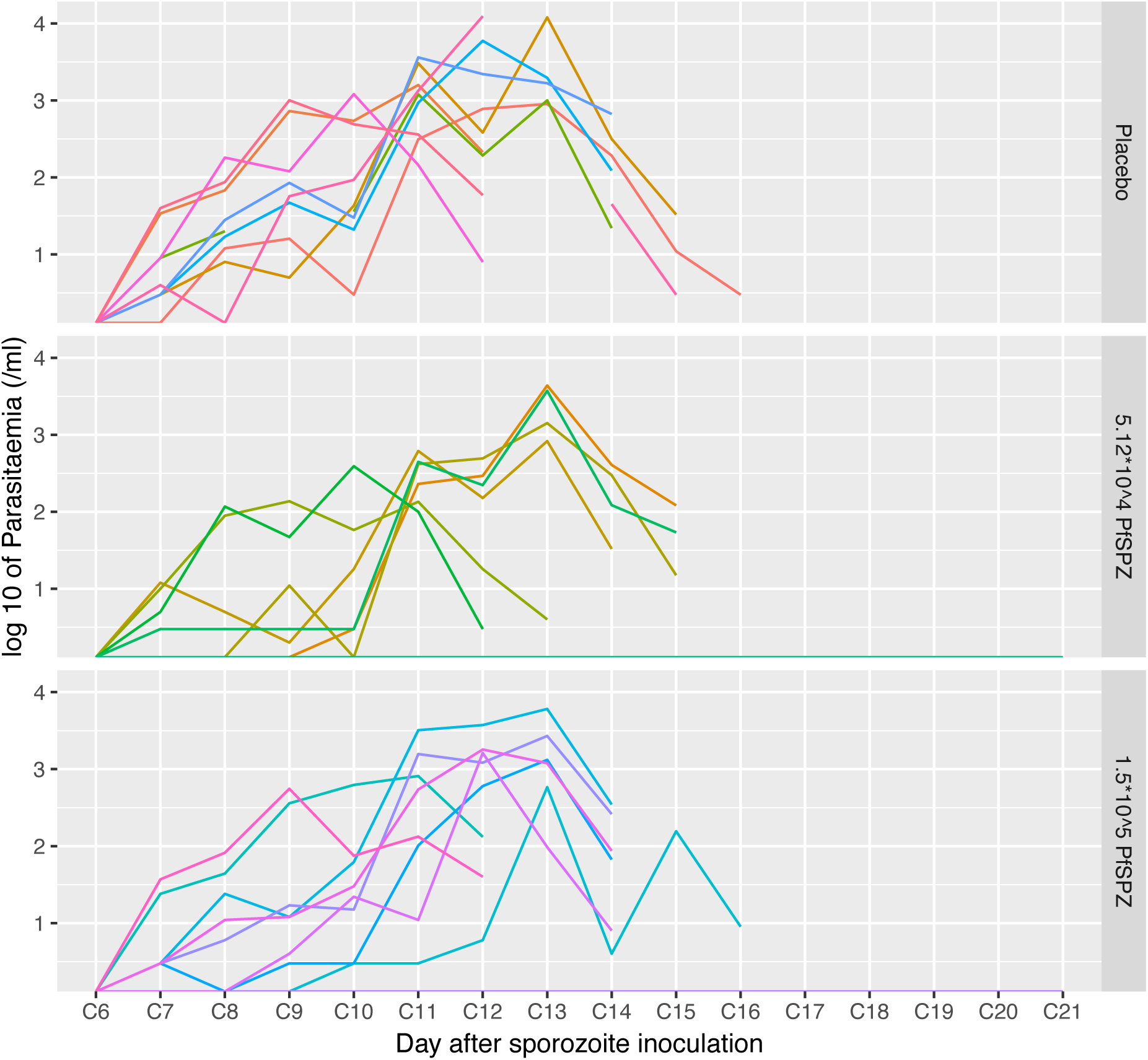
qPCR-assessed Pf parasitemia kinetics in clinical trial participants after CHMI. Shown is the quantification of Pf blood stage parasite load (log10 per ml blood) by qPCR over time after CHMI in the 9 individuals from the placebo control group (top), the 6 non-protected individuals in group A (reference PfSPZ vaccine dose; center) and the 8 non-protected individuals in group B (high PfSPZ vaccine dose; bottom). Curves in different colors depict parasite densities over time in individual participants. Treatment was initiated upon reaching the pre-defined parasitemia endpoint.

**Fig. 4.**
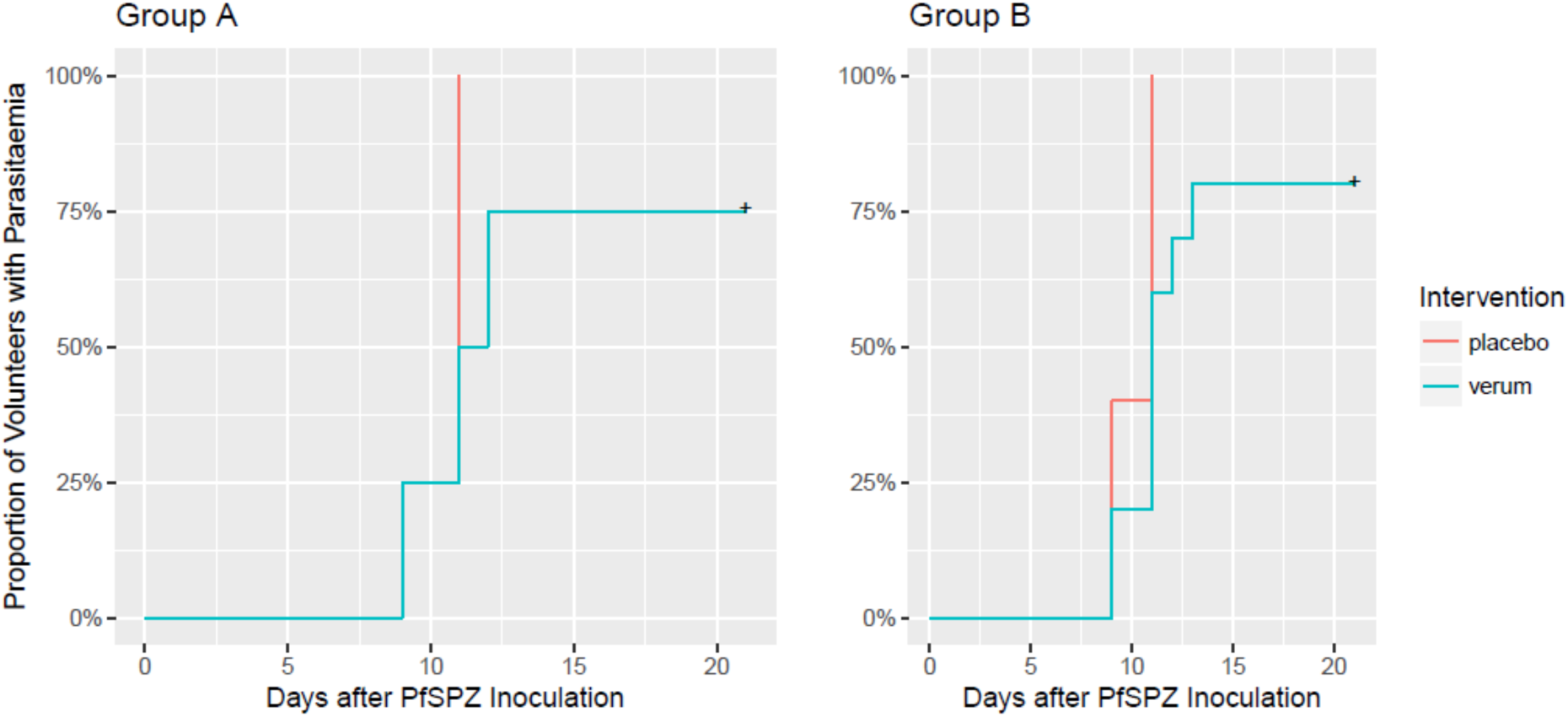
Time to patency after CHMI of participants in groups A and B. Shown are Kaplan-Meier curves of time to initiation of treatment upon reaching a pre-defined parasitemia endpoint. CHMI was done at week 10 after the last immunization in all volunteers, with the exception of one volunteer in Group A (verum) and B (placebo) each, who underwent CHMI at 17 weeks and 14 weeks, respectively. Both were treated for blood infections on day 10 and 12 after CHMI and were included in the graph.

### Anti-PfCSP antibodies generated by early arrest of *Pf*SPZ-CVac immunization

The 25% and 20% VE observed after 3 doses of 5.12x10^4^ or 1.5x10^5^ PfSPZ of PfSPZ-CVac (AP), respectively, contrasts sharply with the 100% VE we achieved with 3 doses of 5.12x10^4^ PfSPZ of PfSPZ-CVac (CQ)(7). To better understand the immunological basis for these differences in VE, we first analyzed the levels of IgG antibodies generated by vaccination against the Pf circumsporozoite protein (PfCSP) two weeks after the 3^rd^ dose of vaccine and just prior to CHMI (Fig. 5). The median serum dilution corresponding to an optical density of 1.0, termed net OD 1.0, two weeks after the last dose of PfSPZ-CVac (AP) was 300 (range, 80 to 4,000) for the 8 subjects who underwent CHMI in vaccine Group A and 3,800 (range 1,400 to 67,100) for the 9 subjects who underwent CHMI in the previous PfSPZ-CVac (CQ) study (7), which used the same dose of 5.12x10^4^ PfSPZ (Table S3)(p < 0.05, Wilcoxon-Mann-Whitney U Test, 2-tailed).

**Fig. 5.**
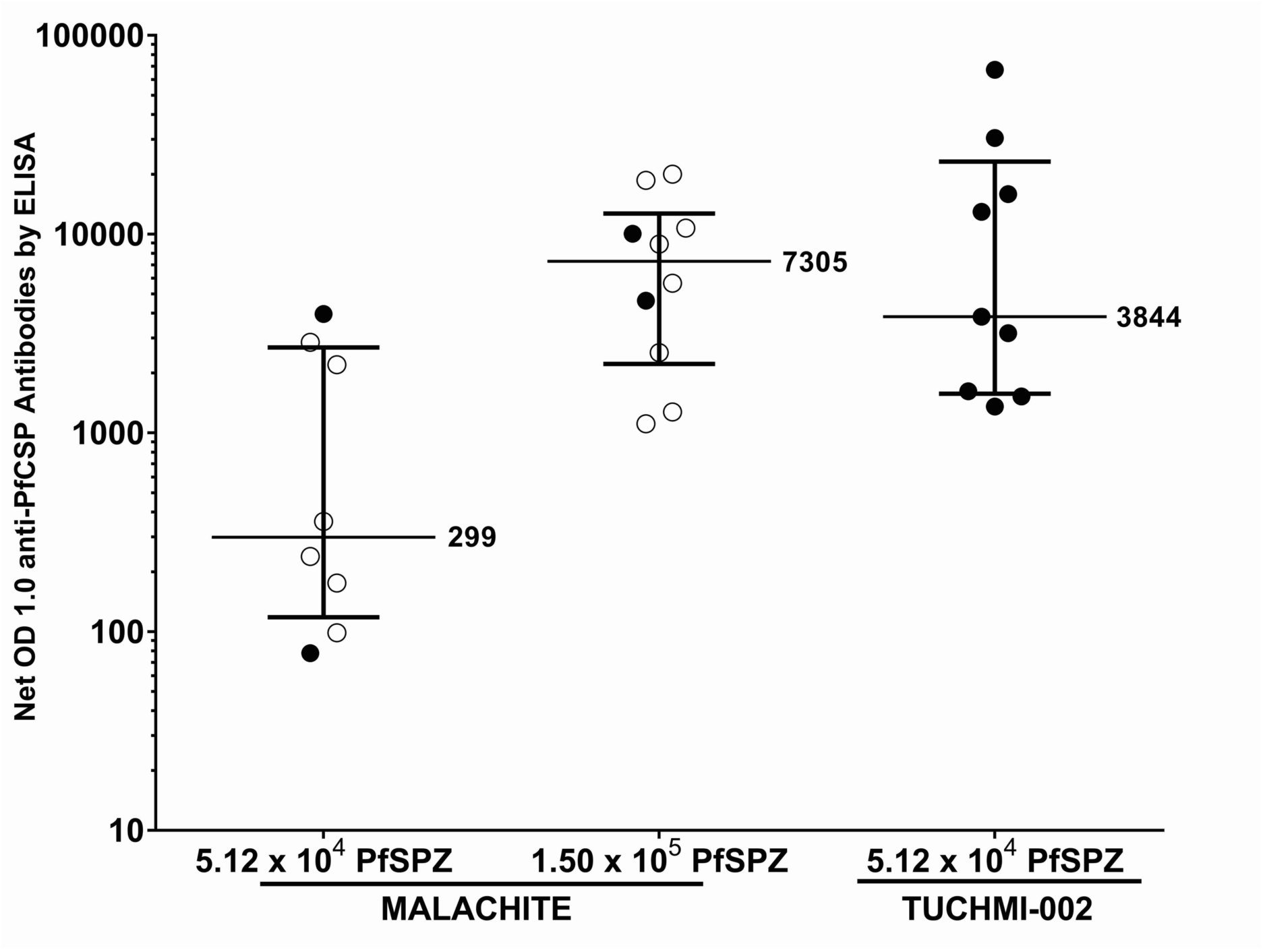
Antibodies to PfCSP 2 weeks after the third dose of vaccine in the current trial (Malachite = PfSPZ-CVac [AP]) compared to data from a previous trial (TÜCHMI-002 = *PfSPZ*-CVac [CQ]) (**7**). Lines represent the median and 25^th^ and 75^th^ quartile levels. Filled circles represent individuals, who did not develop Pf parasitemia (protected), and open circles represent individuals who did develop Pf parasitemia (unprotected). Note that anti-PfCSP antibody levels did not predict VE during CHMI. OD 1.0 is the serum dilution at which the optical density was 1.0. Net OD 1.0 is the OD 1.0 2 weeks after the third dose of vaccine minus the OD 1.0 prior to immunization.

This >10-fold reduction in anti-PfCSP antibody levels in the PfSPZ-CVac (AP) group was unexpected. Since we can firmly exclude batch-to-batch variation during the manufacturing process by GMP-mandated quality control procedures, which includes quantification of sporozoite numbers, motility and invasion capacity, this finding is consistent with a scenario, in which the immune systems of subjects immunized with PfSPZ-CVac (AP) are exposed to less PfCSP than those immunized with PfSPZ-CVac (CQ).

Next we assessed anti-PfCSP antibodies in the higher dose group. Increasing the three PfSPZ doses to 1.5x10^5^ PfSPZ of PfSPZ-CVac (AP) significantly increased median anti-PfCSP antibodies two weeks after the 3^rd^ immunization to 7,300 (range, 1,100 to 20,000, p < 0.01, Mann-Whitney U Test, 2-tailed) (Fig. 5, Table S3). However, despite the > 20-fold increase in PfCSP antibodies in group B there was no increase in VE. We note that the VE of 20% was significantly lower than the 100% VE after immunization with 5.12x10^4^ PfSPZ of PfSPZ-CVac (CQ) (p<0.001, Barnard’s test, 2-tailed). The correlation between antibody concentration prior to CHMI and inoculum were similar (Table S3). Strikingly, the higher levels of anti-PfCSP antibody concentrations in Group B were not predictive of VE after CHMI (Fig. 5). These results are consistent with an anti-PfCSP antibody-independent mechanism of protection after immunization with PfSPZ-CVac (CQ).

### Profiling of IgG antibody responses to sporozoite, liver stage, and asexual blood stage antigens

We hypothesized that reduced protection with PfSPZ-CVac (AP) compared to PfSPZ-CVac (CQ) was due to the early arrest of liver stage development by AP. This pharmaceutical arrest is expected to result in significantly reduced exposure to liver stage and blood antigens. To this end, we probed a representative range of IgG antibody responses with a custom protein microarray (33). We detected a striking absence of a few distinct antibody responses in volunteers who received PfSPZ-CVac (AP) compared to PfSPZ-CVac (CQ) (ref.), while the breadth and intensity of immunoreactivity to 216 other antigens was indistinguishable (Fig. 6A, B). In perfect agreement with the ELISA results (Fig. 5A), IgG responses to *Pf*CSP were considerably higher in volunteers with PfSPZ-CVac (AP) compared to PfSPZ-CVac (CQ) (Fig. 5 and Fig. 6C), likely reflecting higher exposure to sporozoites in the high-dose group of PfSPZ-CVac (AP) (1.5x10^5^ vs 5.12x10^4^, respectively). Most importantly, IgG antibody responses to the two well-known secreted liver stage antigens LISP2 and LSA1 were significantly reduced in recipients of PfSPZ-CVac (AP) (Fig. 5B, C and Supplementary Table S4). Together, our IgG profiling data shows that inferior vaccine efficacy of the PfSPZ-CVac (AP) protocol correlates with low responses to liver stage antigens, whereas superior IgG responses to *Pf*CSP are not predictive of protection.

**Fig. 6.**
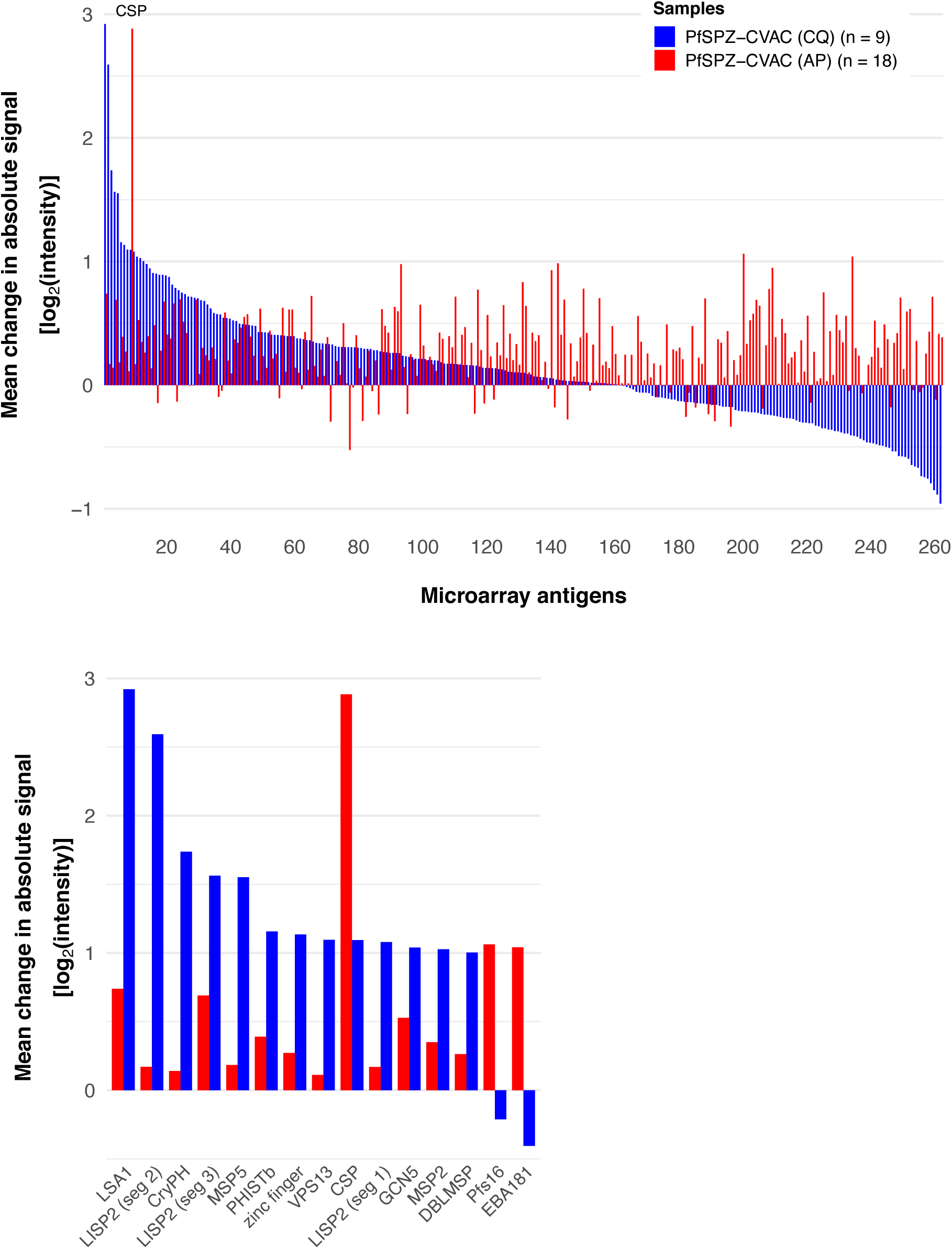
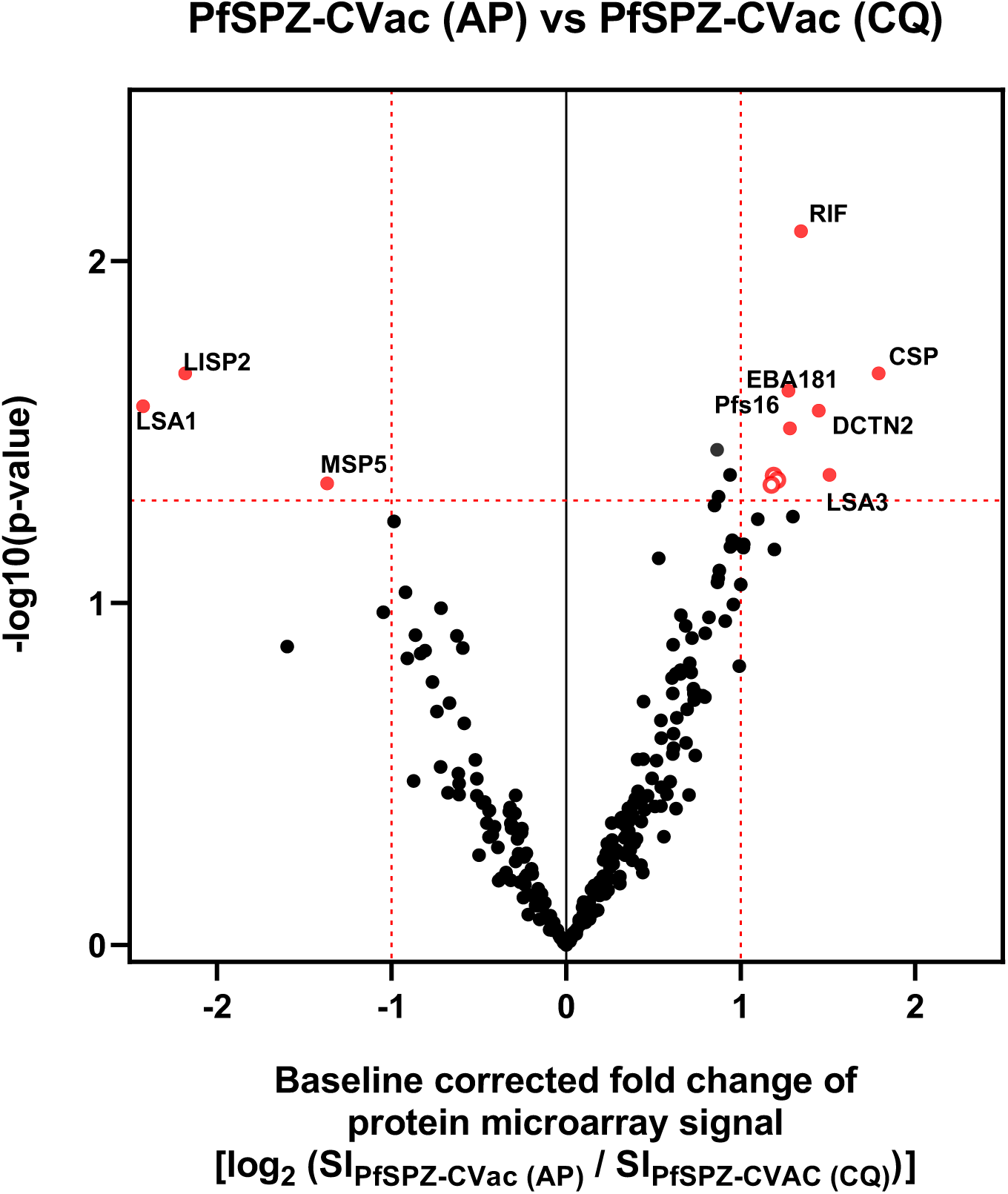
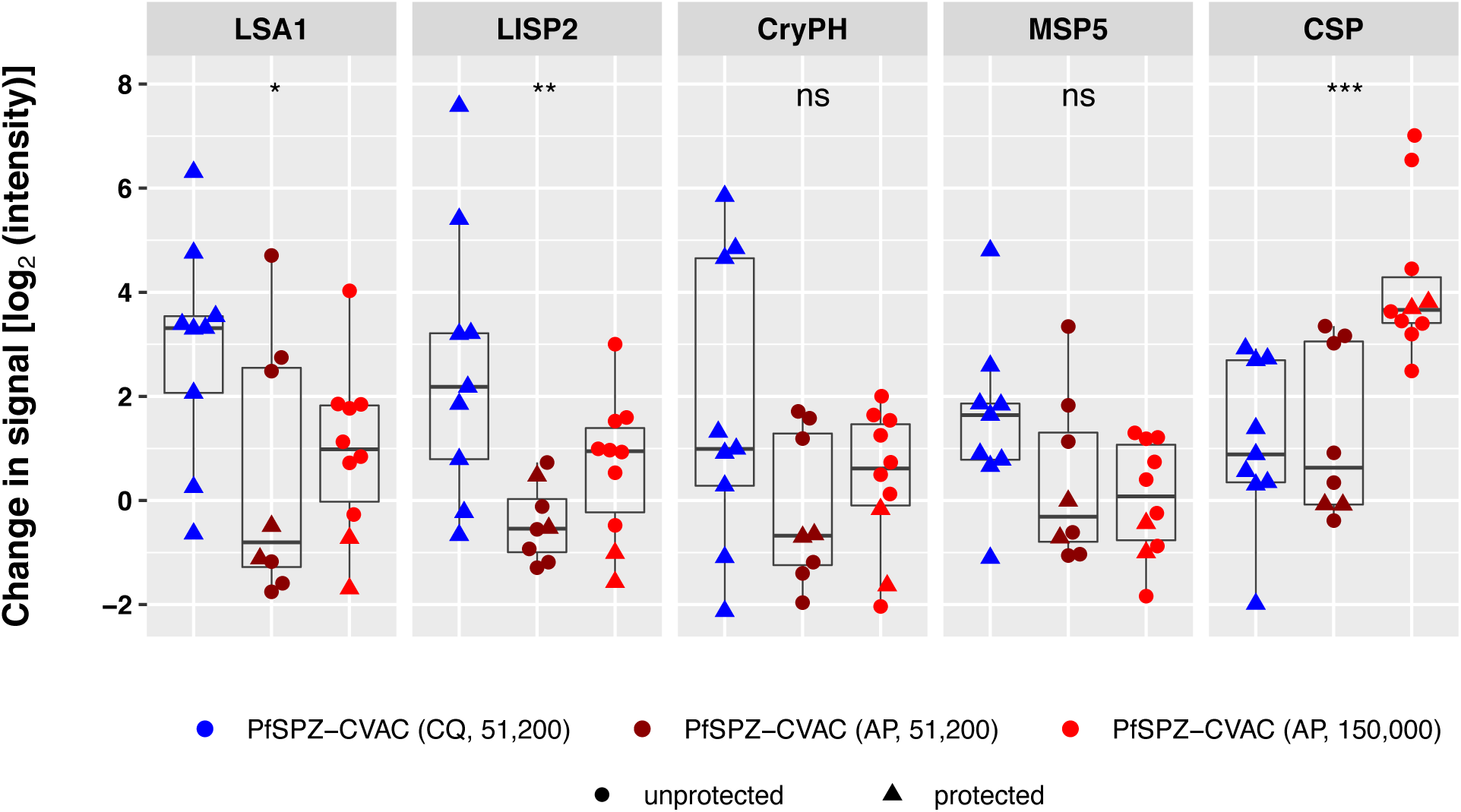
Antibody response measured by protein microarrays comparing reactivities of the two vaccination regimen PfSPZ-CVac (AP) and PfSPZ-CVac (CQ). Sera from all volunteers collected before immunization (baseline, D-1) and one day before challenge (C-1) were applied on protein microarrays at a 1:50 dilution containing 262 Pf proteins representing 228 unique antigens. Analysis was performed on C-1 data after subtraction of the individual baseline reactivity. (A) Bar charts of mean reactivity in the two vaccine regimen, ordered by descending signal intensities for PfSPZ-CVac (CQ) and subset of highest mean changes (> 2-fold change) from both vaccine protocols. Array data were normalized, log2-transformed and baseline reactivity was subtracted. (B) Volcano plot to analyze differential immunoreactivity in the two trials. Antigen reactivity in verum donors of PfSPZ-CVac [AP] (to the right) was compared to the verum donors of PfSPZ-CVac [CQ] (dose 5.12x10^4^ PfSPZ) (to the left). Differentially recognized antigens (p value < 0.05 and fold change > 2) are depicted in red. (C) Box plot of signature IgG responses in PfSPZ-CVac (AP) (stratified by vaccine dose) and PfSPZ-CVac (CQ) (dose 5.12x10^4^ PfSPZ). Asterisks indicate statistical significance.

## Discussion

Intra-host replication is a key feature of many live attenuated vaccines. This expansion correlates with improved protective immunity against natural exposure (34). For instance, the Sabin poliomyelitis vaccine proved that a polio virus strain capable of replicating in the gut but not the nervous system can generate robust neutralizing anti-viral immunity (34). Similarly, it has been proposed that intra-host replication of live, attenuated, PfSPZ-based vaccines might substantially boost the per-parasite VE (5, 7, 16, 35–38).

The current benchmark in humans for highly protective malaria vaccines is immunization with whole PfSPZ, which relies on radiation-, chemo- or genetic attenuation of live, metabolically active PfSPZ (38–41). Immunization with radiation-attenuated (6) and chemo-attenuated (7) PfSPZ vaccines has induced robust and sustained VE in humans against CHMIs for up to 14 months (14, 15) and natural exposure to Pf in the field for at least 6 months (13). Vaccine development has, however, mostly been empirical, and a better understanding of the critical processes involved in the establishment of protective immunity is needed to guide further development of this vaccine strategy.

In Pf, the initial massive (20,000-40,000 fold), but clinically silent, liver stage expansion of parasite biomass and antigenic breadth provides ample opportunities for arresting infections prior to the subsequent pathogenic blood stage. Here, we tested the potential of an attenuation protocol based on AP, a licensed antimalarial drug combination with liver stage activity used for the treatment and chemoprophylaxis of Pf malaria. We proposed that this drug partner would improve the *in vivo* chemo-attenuation strategy significantly because: i) chemoprophylaxis would be administered only concomitantly with PfSPZ increasing safety and practicality, whereas CQ prophylaxis starts prior to PfSPZ injection and is continued with weekly doses until after the last vaccination; ii) there would be no egress of the parasites from the liver, and thus, no possibility of malaria symptoms associated with transient asexual erythrocytic stage parasitemia, which occur with doses beyond 10^5^ PfSPZ; and iii) AP is better tolerated and safer than 4-aminoquinolines, such as CQ.

We demonstrated in the murine malaria model that atovaquone alone and AP led to a complete early arrest of liver stage development. Compared to untreated controls drug-exposed intrahepatic Pb parasites did not begin to replicate as evidenced by single-nucleated liver stage forms observed *in vitro* and by >100-fold reduced parasite liver loads. Of importance, this robust liver arrest was confirmed in the subsequent clinical trial, in which a single dose of AP completely prevented blood stage infections after each of three immunizations with 1.5x10^5^ PfSPZ of PfSPZ Challenge.

Previously, full protection was elicited by three doses of 5.12x10^4^ PfSPZ and chemo-attenuation with CQ (7), which only kills Pf parasites that have emerged from the liver and have commenced subsequent intraerythrocytic replication. In stark contrast, only 2 out of 8 volunteers, who received three doses of 5.12x10^4^ PfSPZ with AP chemoprophylaxis, were protected against CHMI with 3.2x10^3^ *Pf*SPZ of *Pf*SPZ Challenge. Even a 3-fold increase in the PfSPZ numbers per dose to three doses of 1.5x10^5^ PfSPZ failed to induce significant protection despite the robust dose-dependency of PfSPZ-based vaccines (6, 7). Levels of anti-PfCSP antibodies on the day of CHMI in volunteers immunized with three doses of 1.5x10^5^ PfSPZ were at least as high as levels induced by three immunizations with 5.12x10^4^ PfSPZ in the PfSPZ-CVac (CQ) group previously reported (7). In marked contrast, we observed a considerable reduction of IgG antibody responses to the two well-characterized liver stage antigens LISP2 and LSA1. Even though these lower antibody responses may mostly reflect reduced exposure, and not necessarily a protective mechanism, in the PfSPZ-CVac (AP) group compared to the PfSPZ-CVac (CQ) group, it is tempting to speculate that T-cell dependent antibody responses could be a surrogate of cellular immunity. Importantly, unlike our earlier study (7), the comparison of similar PfSPZ doses of PfSPZ-CVac (AP) and PfSPZ-CVac (CQ) allowed us to entangle the effects of sporozoite dose and liver stage maturation, resulting in a clear separation of responses that appear to reflect sporozoite exposure (anti-PfCSP IgG) from responses associated with biological processes critical for protection (anti-LISP2 and anti-LSA1 IgG). Taken together, our data indicate that, high-level VE (≥80%) 10 weeks after the last immunizing dose induced by immunization with 3 doses of 5.12x10^4^ to 1.5x10^5^ infectious PfSPZ under chemoprophylaxis appears to depend on intra-hepatic replication of PfSPZ.

We cannot formally exclude that AP has heretofore undefined immunosuppressive activity in humans as reported for CQ (42). We consider this notion unlikely since over more than 20 years of clinical use of AP no such evidence has been reported. The observed discrepancy between the VE achieved with sporozoite immunization under AP in the murine and human studies (full *vs.* modest protection, respectively) is likely related to the shorter duration of the Pb liver stage (∼2 days) compared to the Pf liver stage (∼7 days). We also noted a largely reduced anti-PfCSP antibody response in) group A vaccinees in comparison to an identical (5.12x10^4^) PfSPZ dose in PfSPZ-CVac (CQ) vaccinees (7). A plausible explanation is that PfCSP is produced and acts as an immunogen not only in extracellular PfSPZ but also throughout Pf liver stage development, coherent with robust CSP expression during liver stage maturation in murine malaria models (43, 44). Accordingly, the full liver stage maturation achieved by PfSPZ-CVac (CQ) immunization as compared to lack of maturation after PfSPZ-CVac (AP) immunization is a likely cause for the marked differences in anti-PfCSP antibody levels after immunization with the same total number of PfSPZ.

We propose that our results indicate an essential role for intrahepatic replication, either through increased amounts of antigen and/or increased numbers of antigens, for achieving maximum protection against CHMI. Furthermore, protection appears to be unrelated to the high anti-PfCSP antibody levels that have been observed previously after high PfSPZ dose immunizations (7). This lack of correlation with antibody responses is supported by data presented here and in previous murine studies showing that the protection induced by radiation-, chemo-, and genetically -attenuated SPZ immunization is strictly dependent on CD8^+^ T-cells (5, 45, 46). Although direct activity of human CD8^+^ T-cells against Pf liver stages in humans has not been demonstrated, our data thus support the hypothesis that cellular immune mechanisms are central to VE of chemo-attenuated PfSPZ vaccines. Further immunological studies are warranted to elucidate the mechanistic basis and identify immune correlates of vaccine-induced protection in humans. It is, of course, conceivable that dependency on intra-host replication can be overcome by administering even higher doses of PfSPZ Challenge with AP, as it has for radiation-attenuated PfSPZ (6, 12, 14, 15), but clinical evaluation of live parasite immunization strategies that combine safe parasite attenuation with superior VE should remain a research priority.

In conclusion, we have demonstrated that generation of full protection against homologous CHMI in humans at an immunizing dose of 5.12x10^4^ to 1.5x10^5^ PfSPZ of PfSPZ Challenge appears to depend on intra-host expansion of Pf, which correlated with an immune signature of anti-liver stage responses. Neither an established rodent malaria model nor anti-PfCSP antibody levels were predictive of sterilizing immunity.

## Methods

### In vitro and animal experiments

All animal work was conducted in accordance with the German Animal Welfare Act (Tierschutzgesetz in der Fassung vom 18. Mai 2006, BGBl. I S. 1207), which implements the directive 86/609/EEC from the European Union and the European Convention for the protection of vertebrate animals used for experimental and other scientific purposes. The protocol was approved by the ethics committee of MPI-IB and the Berlin state authorities (LAGeSo Reg# G0469/09, G0294/15). Female C57BL/6 and NMRI mice at the age of 6 to 8 weeks were purchased from Charles River (Sulzfeld, Germany) for sporozoite injections and blood stage passages, respectively.

### Plasmodium berghei parasites

For all experiments *P. berghei* (Pb) ANKA clone 507, which constitutively expresses the green fluorescent protein (GFP) under control of the *eIF1α* promoter (47), was used. To generate sporozoites, female *Anopheles stephensi* mosquitoes were infected by a bloodmeal on Pb*-* infected NMRI mice. Starting 17 days after infection, salivary glands were hand-dissected from infected mosquitoes, gently ground, and sporozoites harvested after centrifugation. Freshly dissected Pb sporozoites (10^4^) were added to complete DMEM medium (10%FCS, 1% Pen/Strep) containing atovaquone (0.2 µM; Wellvone® Suspension, 750 mg/5ml; GlaxoSmithKline) or Malarone^®^ (0.2 µM atovaquone, 0.08 µM proguanil hydrochloride; GlaxoSmithKline) or the equivalent amounts of DMSO (0.01%) as negative control. Irradiated Pb sporozoites and untreated Pb sporozoites served as control. Sporozoites were added to cultured Huh7 cells in duplicates. Hepatoma cells were incubated for one hour at 37°C for sporozoite sedimentation and for additional 2 hours to permit sporozoite entry. Infected cultures were subsequently washed repeatedly with DMEM complete medium to remove residual drug. Next, infected cells were incubated in complete DMEM medium for 48 h, 69 h or 114 h at 37°C before fixation in 4% paraformaldehyde. Fixed cells were stained with a monoclonal anti-PbHSP70 antibody (48) to visualize parasites, with a rabbit anti-PbUIS4 serum (49) to visualize the parasitophorous vacuole, and with Hoechst 33342 (Invitrogen) that stains nuclei. As secondary antibodies goat Alexa Fluor 488– labeled antibody to mouse immunoglobulin G (IgG) and goat Alexa Fluor 546–labeled antibody to rabbit immunoglobulin G (IgG) (Invitrogen) were used. Image analysis and quantification was performed by fluorescence microscopy using either a Leica DM2500 or a Zeiss Axio Vision microscope. Images were processed with Fiji (Image J, NIH, Bethesda, USA).

### P. berghei sporozoite infection, immunization, and challenge experiments

To test whether single dose administrations of atovaquone or atovaquone-proguanil prevent a subsequent Pb blood stage infection and thus, life-threatening pathology, C57BL/6 mice were intravenously (i.v.) infected with 10^4^ Pb sporozoites, and one dose of drug was co-administered intraperitoneally (i.p.). Drug doses were 3 mg/kg atovaquone alone or 3 mg/kg atovaquone plus 1.2 mg/kg proguanil hydrochloride. Three days later infections were monitored daily by microscopic examination of Giemsa-stained blood films. To test toxicity of atovaquone-proguanil on sporozoites, sporozoites were treated with 1 µM atovaquone-0.4 µM proguanil-hydrochloride in complete DMEM for 2 hours and then washed with RPMI followed by inoculation of 10^4^ treated Pb sporozoites to naïve mice. For quantification of pre-erythrocytic parasite development by qRT-PCR livers were removed 42 hours after sporozoite infection and transferred to TRIZOL^®^ for RNA isolation. Complementary DNA (cDNA) synthesis and qRT-PCR were done as described previously (*4*). Briefly, the mean C_t_ value of the *Pb*18S ribosomal subunit RNA (*18SrRNA*; gene ID: 160641) was normalized to the mean C_t_ of mouse *GAPDH* mRNA values (gene ID: 281199965) using the ΔΔC_t_ method. qPCR experiments were performed with the ABI 7500 sequence detection system and done in triplicates. For immunizations, female C57BL/6 mice were inoculated twice or three times at seven-day intervals with 10^4^ Pb salivary gland sporozoites i.v. together with one dose of drug administered i.p. as described for the chemoprophylaxis studies. Three to five weeks after the last immunization mice were challenged by intravenous injection of 10^4^ Pb salivary gland sporozoites. Naïve, age-matched mice served as infection controls. For determination of sterile protection, blood parasitemia was monitored by microscopic examination of Giemsa-stained blood films daily from day 3 until day 20. Alternatively, the parasite liver load after challenge was quantified by qRT-PCR.

### Anti-sporozoite antibody titers

For titration of *P. berghei*-specific antibodies in mouse serum, salivary gland-associated sporozoites were deposited on glass slides, air-dried and fixed in aceton. After rehydration in PBS and blocking, mouse serum was titrated by serial dilutions and bound antibodies detected with a secondary goat anti-mouse Alexa Fluor 546-coupled antibody (1:1,000). Nuclei were stained with Hoechst 33342 (1:1,000).

### Anti-CSP antibody titers

IgG antibodies to the Pf circumsporozoite protein (PfCSP) were assessed by enzyme linked immunosorbent assay (ELISA) as previously described (7). 96-well plates (Nunc Maxisorp Immuno Plate) were coated overnight at 4 °C with 2.0 µg of the recombinant *P. falciparum* (Pf) circumsporozoite protein (rPfCSP, Lot#122006) protein in 50 µL coating buffer (KPL) per well in coating buffer (KPL). Plates were washed three times with 2 mM imidazole, 160 mM NaCl, 0.02% Tween 20, 0.5 mM EDTA and blocked with 1% Bovine Serum Albumin (BSA) blocking buffer (KPL) containing 1% non-fat dry milk for 1 h at 37 °C. Plates were washed three times and serially diluted serum samples (in triplicate) were added and incubated at 37 °C for 1 h. After three washes, peroxidase labelled goat anti-human IgG (KPL) was added at a dilution of 0.1 µg/mL and incubated at 37 °C for 1 h. Plates were washed three times, ABTS peroxidase substrate was added for plate development, and the plates were incubated for 75 mins at 22°C room temperature. The plates were read with a Spectramax Plus384 microplate reader (Molecular Devices) at 405 nm. The data were collected using Softmax Pro GXP v5 and fit to a 4-parameter logistic curve, to calculate the serum dilution at OD 1.0. A negative control (pooled serum from non-immune individuals from a malaria-free area) was included in all assays. Serum from an individual with anti-PfCSP antibodies for PfCSP was used as positive control.

### Protein microarray and analysis

Microarrays were produced at the University of California Irvine, Irvine, California, USA (33). In total, 262 Pf proteins were expressed using an *E. coli* lysate *in vitro* expression system and spotted on a 16-pad ONCYTE AVID slide, representing 228 important Pf antigens known to frequently appear in sterile and naturally acquired immunity against the parasite (50, 51).

For the detection of binding antibodies, secondary IgG antibody (goat anti-human IgG QDot™800, Grace Bio-Labs #110635), secondary IgM antibody (biotin-SP-conjugated goat anti-human IgM, Jackson ImmunoResearch #109-065-043) and Qdot™585 streptavidin conjugate (Invitrogen #Q10111MP) were used. Study serum samples as well as a European control serum pool were diluted 1:50 in 0.05X Super G Blocking Buffer (Grace Bio-Labs, Inc.) containing 10% *E. coli* lysate (GenScript, Piscataway, NJ) and incubated for 30 minutes on a shaker at room temperature (RT). Meanwhile, microarray slides were rehydrated using 0.05X Super G Blocking buffer at RT. Rehydration buffer was subsequently removed and samples added onto the slides. Arrays were incubated overnight at 4°C on a shaker (180 rpm). Serum samples were removed the following day and microarrays were washed using 1X TBST buffer (Grace Bio-Labs, Inc.). Secondary antibodies were then applied at a dilution of 1:200 and incubated for two hours at RT on the shaker, followed by another washing step and a one-hour incubation in a 1:250 dilution of Qdot™585 streptavidin conjugate. After a final washing step, slides were dried by centrifugation at 500 g for 10 minutes. Slide images were taken using the ArrayCAM® Imaging System (Grace Bio-Labs) and the ArrayCAM 400-S Microarray Imager Software. Microarray data was analyzed in R statistical software package version 3.6.2. All images were manually checked for any noise signal. Each antigen spot signal was corrected for local background reactivity by applying a normal-exponential convolution model (52) using the “saddle” -algorithm for parameter estimation (available in the limma package v3.28.14) (53). Data was log_2_-transformed and further normalized by subtraction of the median signal intensity of mock expression spots on the particular array to correct for background activity of antibodies binding to *E. coli* lysate. Differential antibody levels in the different allocated study outcomes (placebo, non-protected and protected vaccinees) were detected by Student’s t-test. Antigens with p<0.05 and a fold change>2 of mean signal intensities were defined as differentially recognized between the tested sample groups.

### Quantification of antigen-specific CD8^+^ T-cell responses

For CD8^+^ T-cell stimulations followed by intracellular cytokine staining (ICS), splenic lymphocytes were stimulated with 10 µM SSP2/TRAP_130-138_ or S20_318-326_ peptides (24) for 5 hours at 37°C in the presence of brefeldin A (1:1,000). Cells were stained with fluorescently-labeled anti-mouse CD8 [53-6.7], CD62L [MEL14], or CD11a [M17/4] antibodies (eBioscience). Following fixation with 4% paraformaldehyde, cells were stained intracellularly with fluorescently-labeled anti-mouse IFN-γ [R4-6A2] antibody (eBioscience) in permeabilization buffer (BD Bioscience). Antibodies were incubated 60 min at 4°C. After washing and transfer to 1% paraformaldehyde/ PBS cells were quantified using a LSRII flow cytometer (BD Bioscience). Data analysis was performed using FlowJo (Tree Star Inc., Oregon, USA).

### Clinical trial

This trial was approved by the ethics committee of the Eberhard Karls University and the University Hospital Tübingen as well as by the Paul Ehrlich Institute (Langen, Germany) and the Regional Council (Regierungspräsidium Tübingen). The study was compliant with the International Council for Harmonisation Good Clinical Practice guidelines and the German Medicinal Product Act (Arzneimittelgesetz, AMG). The trial was registered at ClinicalTrials.gov (NCT02858817).

This single center, double-blind, randomized, placebo-controlled phase 1 clinical trial was conducted from September 2016 to November 2017 at the Institute of Tropical Medicine in Tübingen, Germany. The study population was selected to represent healthy, malaria-naïve adults. Volunteers aged 18–45 years from Tübingen and surrounding area with a body-mass index (BMI) between 18 kg/m² and 30 kg/m² were included. Female participants were required to practice effective contraception and to provide a negative pregnancy test. Further inclusion criteria included: being reachable at all times by mobile phone during the whole study, agreement to share medical information about the volunteer with his or her general practitioner, and understanding of study procedures and risks, assessed by a quiz. Additionally, willingness to undergo CHMI with PfSPZ Challenge, to take a curative regimen of antimalarial if necessary, and the ability to comply with all study requirements (in the investigator’s opinion) were also required.

Exclusion criteria were: a history of malaria or plans to travel to endemic regions during the study, receiving any investigational product in another clinical trial within 90 days before enrolment or planned receipt during the study, previous participation in a malaria vaccine trial, history of serious psychiatric conditions, convulsions, or severe head trauma, any malignancy, and diabetes mellitus. Moreover, falling in moderate risk or higher categories for fatal or non-fatal cardiovascular event within 5 years (54), prolonged QTc interval (>450 ms), or any other clinically significant abnormalities in the electrocardiogram, breast feeding, or intention to become pregnant, HIV, hepatitis B or C virus infection, alcohol or drug abuse, any suspected immunodeficient state, history of splenectomy, and haemoglobinopathies also prevented participation. A complete list of eligibility criteria is available as an online supplement. Eligibility criteria were assessed after written informed consent was given.

Fifteen volunteers per group were enrolled and randomized to receive Sanaria® PfSPZ Challenge for immunization or normal saline placebo with an allocation ratio of 2:1 for vaccine:placebo. In Group A, participants received 5.12x10^4^ PfSPZ of PfSPZ Challenge (NF54) by DVI three times at four week intervals and a single dose of AP (1,000 mg/400 mg) administered orally within one hour before each immunization. In Group B participants received 1.5x10^5^ PfSPZ of PfSPZ Challenge (NF54) by DVI with the same scheduling and chemoprophylactic regimen.

Ten weeks after the last immunization, the first CHMI was performed in both groups for vaccine efficacy (VE) testing. CHMI was done by DVI of 3.2x10^3^ PfSPZ of PfSPZ Challenge (NF54). Active follow-up of the participants was conducted through 56 days after the injection of PfSPZ Challenge for CHMI. The protocol stipulated two successive CHMIs, the first at 10 weeks and the second at 16-44 weeks. The sequence of CHMI was planned to be PfSPZ Challenge (NF54, homologous clone) followed by PfSPZ Challenge (7G8, heterologous clone) for Group A. The sequence for Group B was to be based on VE following first CHMI in Group A: NF54 followed by 7G8 when VE against homologous CHMI was <75%, 7G8 followed by 7G8 when VE was ≥75%. Due to the low efficacy of the first CHMI also in group B a second CHMI was not performed in Group B. First CHMI was thus performed with PfSPZ Challenge (NF54) for both groups.

PfSPZ Challenge (NF54) is comprised of aseptic, purified, cryopreserved NF54 PfSPZ, produced by Sanaria Inc. (Rockville, US). PfSPZ Challenge was stored and transported in liquid nitrogen vapor phase at -150 to -196°C. Formulation and reconstitution was made in Tübingen on the day of infection. Volunteers were inoculated within 30 minutes after thawing of PfSPZ Challenge. Sterile isotonic normal saline, identical in appearance to PfSPZ Challenge was used as placebo. A volume of 0.5 mL of vaccine or placebo was injected into an arm vein by DVI through a 25 gauge needle. After each immunization, participants were monitored for at least 60 minutes before leaving the clinic for local and systemic adverse events. Participants were assessed on site for safety and to measure parasitemia on days 1, 5, 7, 10, 14 and 21 after each immunization, and on day 1, 6-21, 28, 56 after CHMI. Medically qualified study personnel were available continually for unscheduled visits. Antimalarial treatment, according to the German guidelines (55) for the treatment of uncomplicated Pf malaria (AP or artemether-lumefantrine as first-line drugs) was to be initiated, in the event of breakthrough parasitemia with symptoms during immunization or in the case of parasitemia following CHMI. Breakthrough parasitemia was defined as microscopically detectable parasitemia during immunization with at least two symptoms consistent with malaria for 2 days despite chemoprophylaxis. Protection was defined as the absence of parasites in the peripheral blood for 28 days following CHMI. Parasitemia after CHMI was assessed on daily basis from day 6 to day 21 and again on day 28 via thick blood smear (TBS) and quantitative PCR. Treatment was administered upon occurrence of three consecutive positive PCR results one of them at least 100 Pf parasites/ml from samples taken at least 12 hours apart or the first TBS positivity. An additional follow-up visit for safety was conducted on day 56. Participants were encouraged to immediately report adverse events between the scheduled follow-up visits.

If a volunteer was withdrawn from the study after receiving a dose of PfSPZ Challenge at one or more of the three immunizations or at CHMI, a full, appropriate, curative course of antimalarial therapy was administered.

In both groups 10 volunteers were randomly allocated to receive immunizations with PfSPZ Challenge and 5 volunteers to receive normal saline placebo (PfSPZ Challenge (NF54):placebo = 2:1). Group membership was allocated using a Mersenne-Twister random number generator implemented in R. A third party outside the study team generated and distributed the randomization list. A dedicated member of the formulation team, who was not involved in volunteer management or diagnostic activities, kept the randomization envelopes and dosing schedule.

The primary aim of the study was to assess the safety and VE of repeated immunization by DVI of PfSPZ Challenge under AP chemoprophylaxis in malaria naïve adults. The primary VE endpoint was the proportion of protected volunteers.

Protection was defined as the absence of parasites in the peripheral blood for 28 days following first CHMI with PfSPZ Challenge. To assess safety outcomes, Grade 3 and 4 adverse events (AEs) and serious adverse events (SAEs) were captured from time of first administration of A/P until the end of the study. Functional characterization of humoral and cellular immune responses were exploratory endpoints.

### Statistical analysis and power calculation of clinical trial

To be able to show, with a power of 80% and a two-tailed alpha of 5%, that 25% or less of immunized volunteers and 95% of controls, allocated in a 2:1 ratio became infected by CHMI, 10 immunized and 5 placebo-treated volunteers per group were required. Hence, a total of 30 (10 each for 5.12x10^4^ and 1.5x10^5^ *Pf*SPZ Challenge (NF54) with AP and 10 placebo) volunteers were required. The sample size was calculated using the nBinomial function in the gsDesign package.

No formal hypothesis testing was done for safety and tolerability data. Safety and tolerability data are presented as descriptive analyses in listings and graphically. VE was calculated by comparison of proportions between immunized and placebo-treated volunteers using an unconditional exact test (Boschloo’s test) and time-to-parasitemia using a log-rank (LR) test. Multiple non-parametric group comparisons were performed by using the Kruskal-Wallis (KW) H test. The level of significance was set at a two-tailed type 1 error alpha <5%. All statistical analyses were performed using R version 3.4.4 and GraphPad Prism 5. Kaplan-Meier curves were compared by a log-rank (Mantel Cox) test. Statistical significance of parasite sizes, qPCR data, and flow cytometric analyses were assessed using the Mann-Whitney U test for nonparametric test samples. *P* values of *P* < 0.05 were considered as significant.

## Acknowledgments

Funding for the clinical trial was provided by the German Centre for Infection Research (DZIF). We are grateful for the support provided in part by National Institutes of Allergy and Infectious Diseases, National Institutes of Health (USA), SBIR grants 5R44AI055229 and 2R44AI058375 to SLH. The preclinical work was supported by a DFG grant (SFB 544) to SB and KM and by the Max Planck Society to KM. The clinical trial was funded by AKF grant 346-0-0.

We wish to acknowledge the volunteers participating in the clinical trial. We also gratefully acknowledge the contribution of the manufacturing, quality, regulatory and clinical teams at Sanaria.

Authors listed as Sanaria affiliates are all full time employees of Sanaria Inc., Rockville MD.

Author contributions:

Design: SB, SLH, TLR, PGK, KMa, BM

Principal Investigator and clinical trial sponsor representative: PGK

Writing: SB, MS, KMü, ZS, RF, FOR, TLR, PGK, SLH, KMa, BM

Preclinical experiments: KMü, JF, JH

Clinical trial: MS, ZS, AL, TLS, TTN, JI, HLH, DMW, RS, SA, PGB, ZM, ME, WM, TG, FOR, JH GMP: ERJ, AR, YA, SC, AM, NKC, PB, BKLS

Pharmaceutical Operations: ERJ, AR, YA, AM, NKC, BKLS, CLC, AK

Immunology: KMü, JH, NKC, SC, RF, FRL

