## Supplementary Material for "Mapping of safe and early chemo-attenuated live *Plasmodium falciparum* immunization identifies immune signature of vaccine efficacy"

**Content:**

- **Supplemental Information**
- **Supplemental Figures S1 - S5**
- **Supplementary Tables S1 - S4**
- **Supplemental References**

**Supplemental information: Safety of immunization with PfSPZ-CVac with AP chemoprophylaxis**

The overall frequencies of grade 1-4 adverse events were similar in recipients of PfSPZ-CVac and normal saline placebo (487 in 20 verum volunteers vs. 221 in 10 placebo volunteers). Of the overall 708 AEs, 483 were considered unrelated or unlikely to be related, 91 as possibly related, 124 as probably related and 8 as definitely related to the administration of PfSPZ Challenge (summarized in Table 2). The most frequent AE was headache (n=81), followed by increase in diastolic blood pressure (n=61) and systolic blood pressure (n=79). One serious adverse event occurred after the first vaccination: a female patient was hospitalized 17 days after the first vaccination with left lower abdominal pain and diagnosed with an ovarian cyst rupture. This SAE was deemed to be unrelated to the study. In total 531 grade 1, 132 grade 2, 40, grade 3 and 5 grade 4 AEs were recorded. All grade 4 AEs (including the above described SAE with ovarian cyst rupture and associated abdominal pain, hypoglycemia and two episodes of creatinine kinase elevation) were considered as unrelated or unlikely to be related to the study treatment. The mild local and systemic symptoms after immunization (tenderness, pruritus of the injection site, headache, fatigue, dizziness, laboratory changes) were transient and resolved within a few days.

Of the 27 individuals who underwent CHMI, 23 developed Pf parasitemia. 21 of these 23 experienced at least one mild or moderate (grade 1 or 2) symptom associated with Pf parasitemia or antimalarial treatment. Four related grade 3 AEs occurred: lymphocytopenia, thrombocytopenia, fever, subcostal pain. All study-associated AEs resolved completely without sequelae.

### Supplementary Figure S1

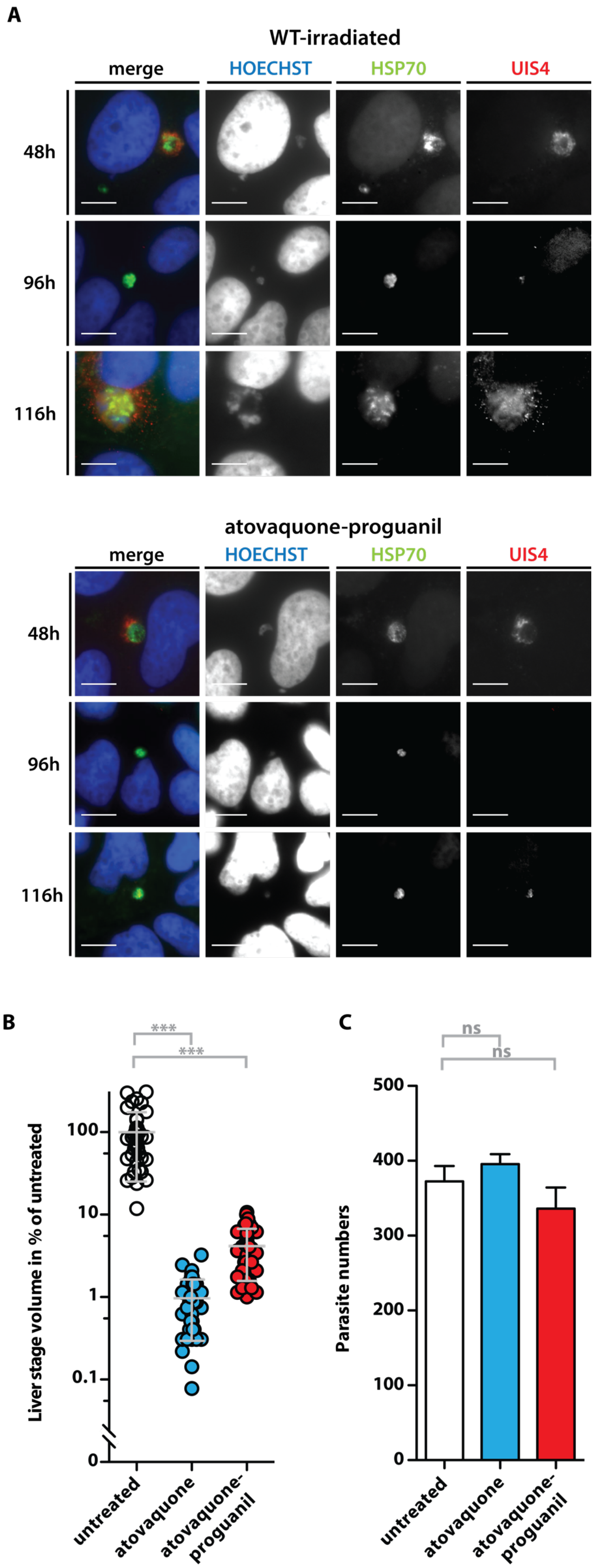

**Fig S1. Pb liver stages treated with atovaquone-proguanil retain their invasive capacity, are fully arrested, and persist 5 days after infection *in vitro*.**

(A) Composite fluorescence micrographs of mature *Plasmodium berghei* liver stages in cultured hepatoma cells. Shown are representative images of liver stages 48h, 96h and 116h after infection with sporozoites. During the first three hours cultures were exposed to atovaquone-proguanil. Irradiated, untreated sporozoites served as a control. Parasites were visualized by fluorescent staining of the cytoplasm (green; anti-PbHSP70 antibody), the parasitophorous vacuolar membrane (red; anti-PbUIS4 anti-serum), and nuclei (blue; Hoechst 33342). Scale bars: 10 $\mu$m.

(B) Quantification of liver stage volumes after prophylactic drug treatment. Parasite volume was quantified 48 hours after infection and normalized to the average volume of untreated parasites. Shown are mean percentages (±S.D.). ***, $p<0.001$ (Mann-Whitney U test).

(C) Sporozoite invasion is unaffected by short-term drug treatment. Shown are mean numbers of liver stages (±S.D.) in cultured hepatoma cells 48 hours after infection with untreated, atovaquone-treated or atovaquone-proguanil-treated sporozoites. *ns*, non-significant (Mann-Whitney U test).

**Supplementary Figure S2**

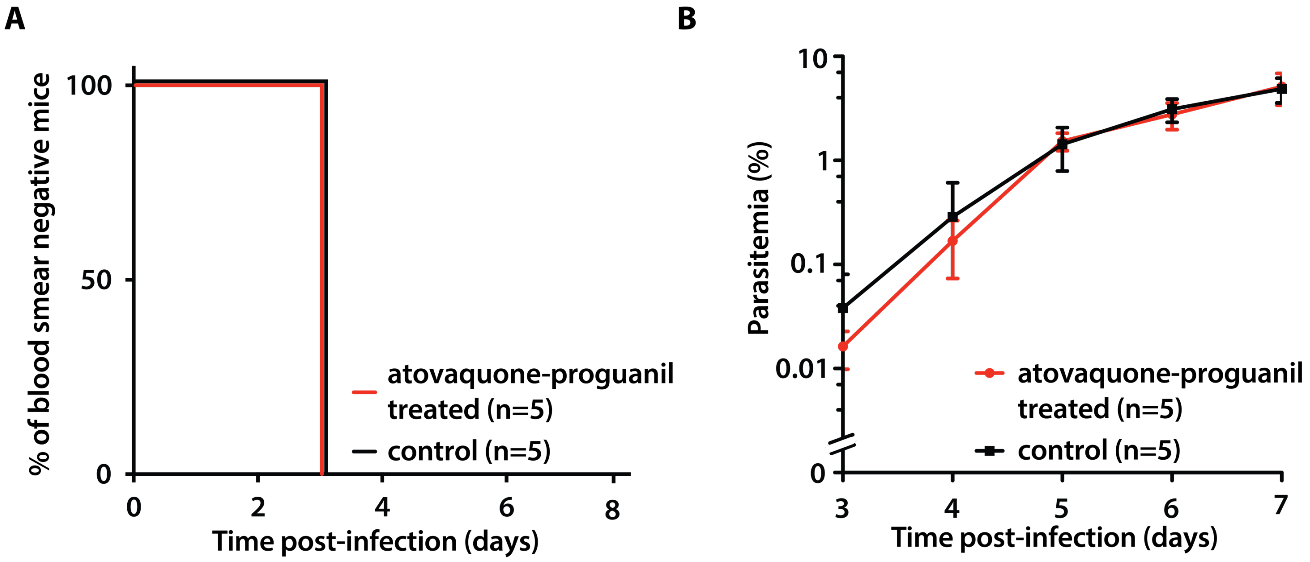


**Fig S2. Chemo-attenuation with atovaquone-proguanil does not interfere with assessment of protection.**

(A) Atovaquone-proguanil does not inhibit infections in naïve mice when challenged 3 weeks after drug administration. Kaplan Meier analysis of time to blood infection upon challenge of mice with $10^4$ sporozoites that previously received atovaquone-proguanil only, given at intervals of the immunization protocol. C57BL/6 mice were treated three times at weekly intervals with 3/1.2 mg/kg atovaquone-proguanil (red line) i.p. or left untreated (black line).

(B) Kinetics of blood stage infections after challenge of mice with sporozoites as described in (A). Parasitemia was determined by daily microscopic examination of Giemsa-stained blood films. Shown are mean asexual blood stage parasite densities (±S.D.).

### Supplementary Figure S3

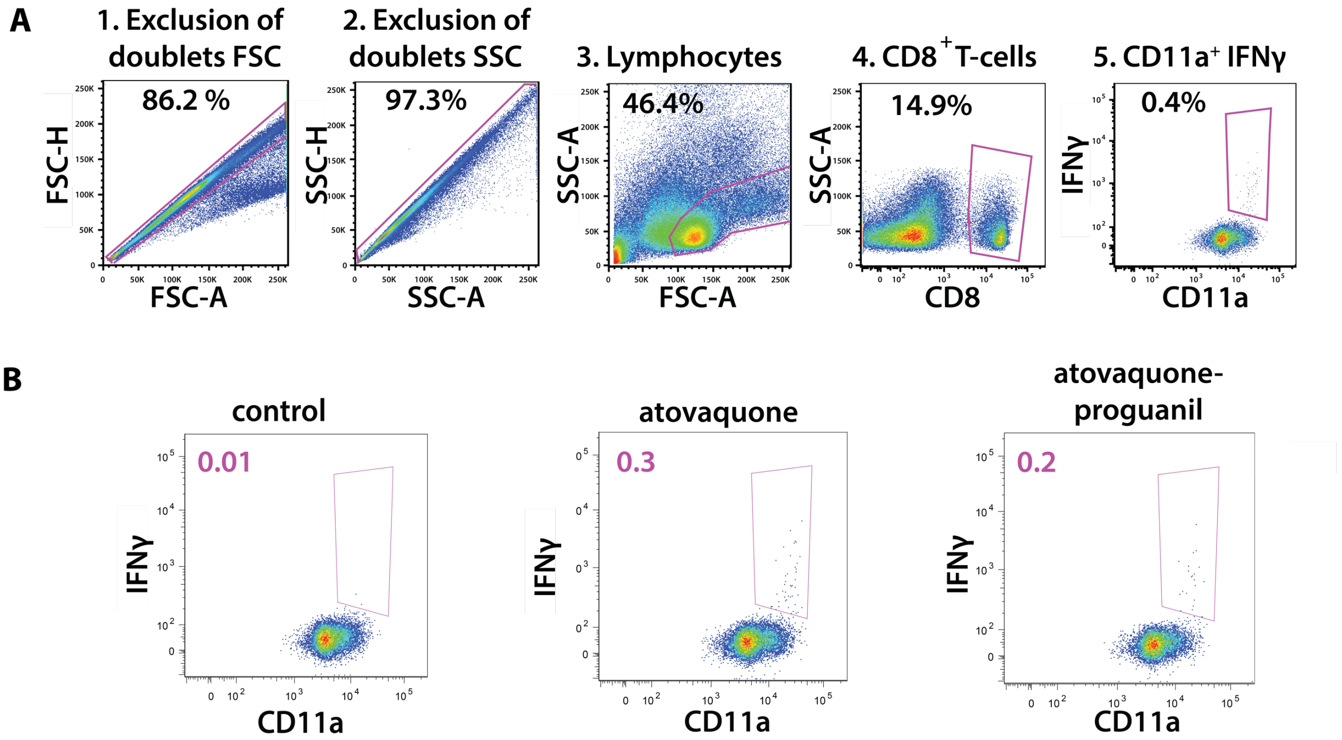


**Fig S3. Identification of IFNγ-secreting activated CD8$^+$ CD11a$^+$ T-cells after re-stimulation with the SSP2/TRAP$_{130-138}$ and S20$_{318-326}$ peptides.**

(A) Shown are exemplary FACS plots illustrating the gating strategy for IFNγ secretion by CD8$^+$ CD11a$^+$ T-cells after re-stimulation.

(B) Representative FACS plots of CD8$^+$ CD11a$^+$ T-cells positive for intracellular IFNγ expression after re-stimulation with the SSP2/TRAP$_{130-138}$ peptide. Numbers show percentage of IFNγ produced by CD8$^+$ CD11a$^+$ T-cells.

**Supplementary Figure S4**

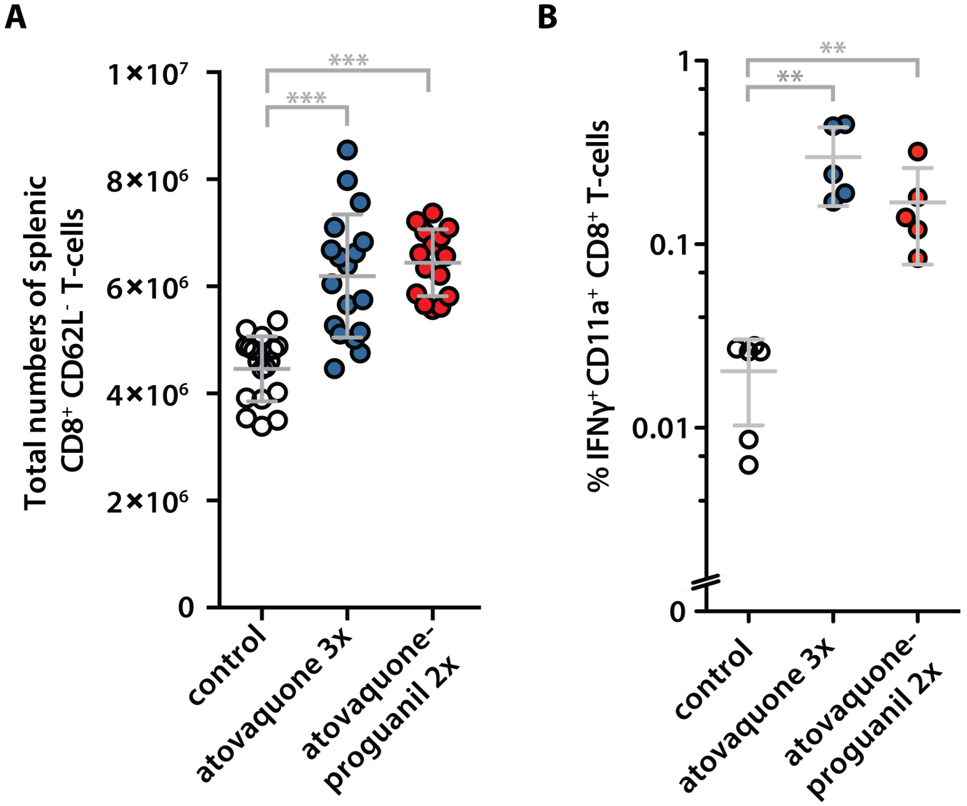


**Fig S4. Total numbers of effector memory T-cells and proportion of $S20_{318\text{-}326}$- experienced CD8+ CD11a+ T-cells.**

Mice (n≥5 each) were immunized by co-administration of sporozoites and a single dose of atovaquone (3 mg/kg i.p.) or atovaquone-proguanil (3/1.2 mg/kg i.p.). Mice were immunized twice (atovaquone-proguanil co-administration, red circle, n=5) and three times (atovaquone co-administration, blue circle, n=6). Naïve mice served as controls (white circle; *n*=6). Sporozoite challenge was done by I.V. injection of $10^4$ sporozoites three to four weeks after the last immunization.

(A) Quantification of total $CD8^+$ $CD62L^-$ T-cells from spleens of immunized or control mice (measured in triplicates). Shown are mean values (±S.D.). ***, $p < 0.001$ (Mann-Whitney U Test).

(B) Quantification of $S20_{318\text{-}326}$ peptide-specific IFNγ-secretion by $CD8^+$ $CD11a^+$ T-cells from spleens of immunized or control mice. Shown are mean values (±S.D.). **, $p < 0.01$(Mann-Whitney U Test).

**Supplementary Figure S5**

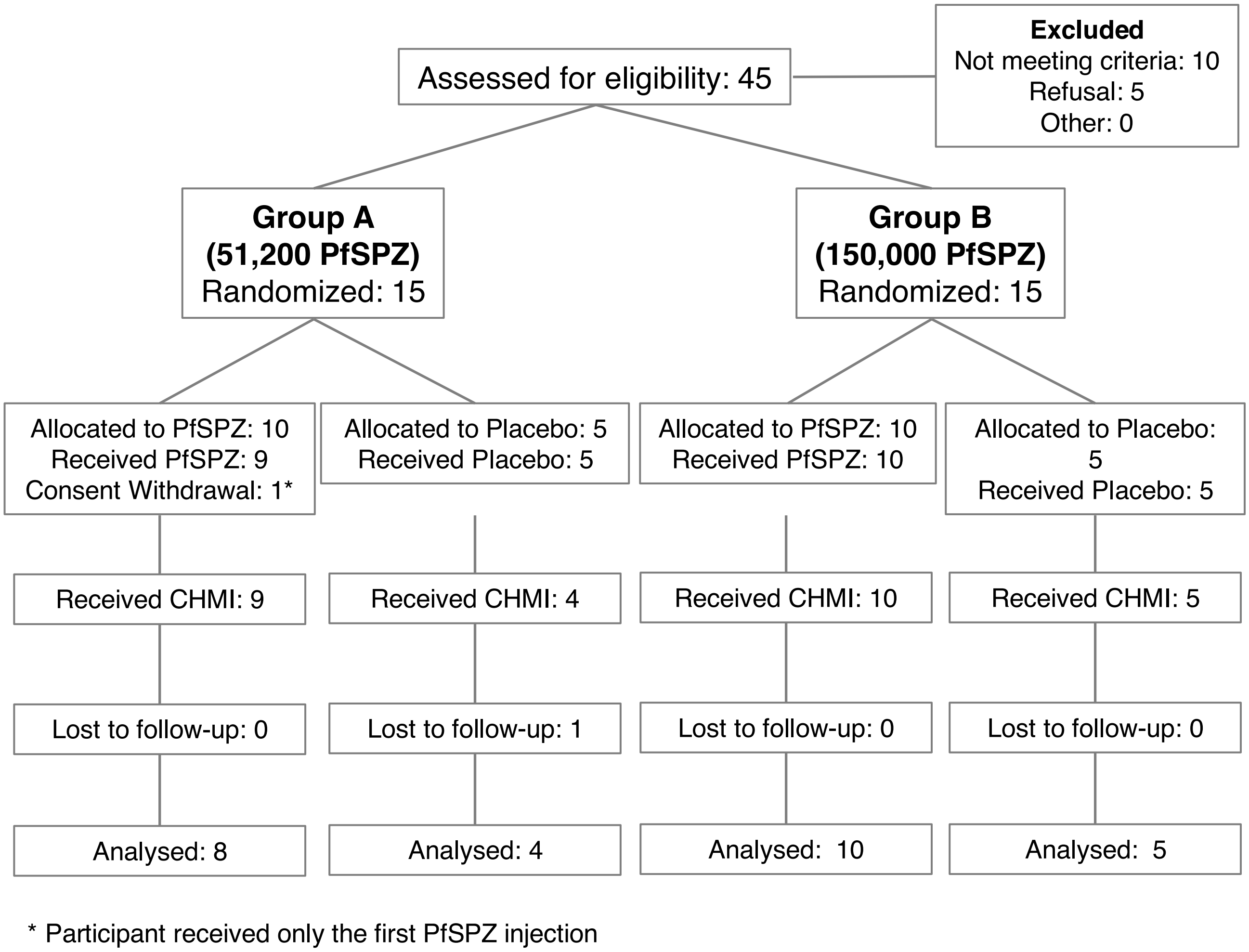


**Fig. S5. Study flow chart**

**Supplementary Table S1. Baseline characteristics of study population.**

| | Vaccinees | | Placebo |
|---|---|---|---|
| | **$5.12x10^4$ PfSPZ** | **$1.5x10^5$ PfSPZ** | |
| | n=10 | n=10 | n=10 |
| **Sex** | | | |
| Female | 7 | 6 | 5 |
| Male | 3 | 4 | 5 |
| **Age (years)** | | | |
| Mean±SD | 26.9 ±6.8 | 28.1 ±2.2 | 24.7 ±4.6 |
| Median | 24 | 28 | 24 |
| Min, Max | 19, 36 | 26, 32 | 19, 32 |
| **BMI** | | | |
| under 18.5 | 1 | 1 | 1 |
| 18.5-24.9 | 7 | 5 | 6 |
| 25.0-29.9 | 2 | 4 | 3 |
| 30.0 or over | 0 | 0 | 0 |
| Mean±SD | 22.4 ±3.5 | 22.2 ±3.2 | 24 ±3.4 |
| Min, Max | 18.1, 28 | 18.1, 27 | 18.3, 29 |

**Supplementary Table S2. Most frequent related adverse events.**

| | Placebo | Vaccinees | |
|---|---|---|---|
| | | **5.12x10^4 PfSPZ** | **1.5x10^5 PfSPZ** |
| | n=10 | n=10 | n=10 |
| **AEs related to immunization** | **33** | **19** | **13** |
| | Headache 21% (7)<br>• grade 1 (6)<br>• grade 2 (1)<br>Dizziness 12% (4)<br>• grade 1 (4)<br>Fatigue 5% (3)<br>• grade 1(3)<br>Tachycardia 5% (3)<br>• grade 1 (3)<br>Miscellaneous (48%) | Hypertension 31% (6)<br>• grade 1 (3)<br>• grade 2 (2)<br>• grade 3 (1)<br>Dizziness 21% (4)<br>• grade 1 (4)<br>Pruritus at injection site 10.5% (2)<br>• grade 1 (2)<br>Headache 10.5% (2)<br>• grade 1 (1)<br>• grade 2(1)<br>Diarrhea 10.5% (2)<br>• grade 1 (2)<br>Miscellaneous (16%) | Fatigue 23% (3)<br>• grade 1 (2)<br>• grade 2 (1)<br>Neutropenia 30% (4)<br>• grade 1 (3)<br>• grade 2 (1)<br>Miscellaneous (46%) |

| | | | |
|---|---|---|---|
| **AEs related to CHMI** | **75** | **44** | **37** |
| | Lymphopenia 13% (10)<br>• grade 1 (5)<br>• grade 2 (4)<br>• grade 3 (1)<br>Fever/Feverish 9% (7)<br>• grade 1 (2)<br>• grade 2 (4)<br>• grade 3 (1)<br>Headache 12% (9)<br>• grade 1 (6)<br>• grade 2 (3)<br>Fatigue 9% (7)<br>• grade 1 (5)<br>• grade 2 (2)<br>Nausea 5% (4)<br>• grade 1 (4)<br>Sweating 5% (4)<br>• grade 1 (3)<br>• grade 2 (1)<br>Miscellaneous (54.6%) | Headache 16% (7)<br>• grade 1 (5)<br>• grade 2 (2)<br>Lymphopenia 9% (4)<br>• grade 1 (2)<br>• grade 2 (2)<br>Fatigue 9% (4)<br>• grade 1 (3)<br>• grade 2 (1)<br>Diarrhea 7% (3)<br>• grade 1 (3)<br>Miscellaneous (59%) | Headache 16% (6)<br>• grade 1 (4)<br>• grade 2 (2)<br>Neutropenia 16% (6)<br>• grade 1 (4)<br>• grade 2 (2)<br>Fatigue 16% (6)<br>• grade 1 (5)<br>• grade 2 (1)<br>Lymphopenia 13% (5)<br>• grade 1 (3)<br>• grade 2 (2)<br>Miscellaneous (38%) |

**Supplementary Table S3. Antibodies to *Pf*CSP two weeks after the third dose of vaccine and 8 weeks (pre-CHMI) after the third dose of vaccine.**

| Group | Volunteer ID | Net OD 1.0 *Pf*CSP ELISA | |
|---|---|---|---|
| | | 14 d Post 3rd Dose | Pre-CHMI |
| **Group A 5.12x10$^4$ *Pf*SPZ (AP)** | MT03 | 2204 | 2027 |
| | MT07 | 99 | 65 |
| | MT08 | 2861 | 1918 |
| | MT10 | 359 | 326 |
| | MT14 | 3967 | 21,836 |
| | MT15 | 239 | 426 |
| | MT16 | 176 | 88 |
| | MT20 | 78 | 134 |
| | **Median** | **299** | **376** |
| **Group B 1.5x10$^5$ *Pf*SPZ (AP)** | MT21 | 10,067 | 13,421 |
| | MT22 | 1276 | 2076 |
| | MT23 | 10,744 | 8840 |
| | MT26 | 18,649 | 18,870 |
| | MT30 | 20,039 | 4823 |
| | MT32 | 5676 | 2424 |
| | MT35 | 4633 | 15,050 |
| | MT36 | 1113 | 835 |
| | MT37 | 8933 | 4542 |
| | MT40 | 2540 | 2306 |
| | **Median** | **7305** | **4683** |

| | | | |
|---|---|---|---|
| **Control (AP)** | MT01 | -13 | 3 |
| | MT02 | 8 | 2 |
| | MT05 | -13 | ND |
| | MT13 | -10 | 3 |
| | MT28 | 18 | -2 |
| | MT31 | 10 | -17 |
| | MT39 | 5 | 37 |
| | MT42 | ND | ND |
| | MT53 | 32 | -7 |
| | **Median** | **7** | **2** |
| **TUCHMI-002 5.12 x $10^4$ PfSPZ (CQ)** | 017 | 1621 | 849 |
| | 026 | 1357 | 466 |
| | 035 | 1529 | 632 |
| | 042 | 3844 | 1536 |
| | 051 | 15,914 | 9056 |
| | 052 | 3180 | 2025 |
| | 071 | 30,518 | 11,503 |
| | 072 | 67,088 | 34,651 |
| | 073 | 12,964 | 5493 |
| | **Median** | **3844** | **2025** |

*Antibody levels are reported as net OD 1.0*, which is the OD 1.0 at the specified time point minus the OD 1.0 prior to immunization.

*OD 1.0 = serum dilution at which the optical density was 1.0.

**Table S4. Differentially recognized antigens, expression profiles and inclusion into subunit vaccine strategies**

| | | | | mRNA expression [1-6] | | | Protein expression [4-8] | | | |
|---|---|---|---|---|---|---|---|---|---|---|
| Protein | Gene ID | Description | Surface | SPZ | LS | BS | SPZ | LS | BS | Subunit vaccine |
| Liver stage antigen 1 (LSA1) | PF3D7_1036400 | dominant liver stage antigen | - | - | - | - | - | + [9] | - | yes [10] |
| Liver specific protein 2 (LISP2) | PF3D7_0405300 | exported to hepatocyte | - | - | + | - | - | + | - | - |
| Merozoite surface protein 5 (MSP5) | PF3D7_0206900 | duplicated antigen | yes | + | - | + | + | - | + | yes [11] |
| Circumsporozoite protein (CSP) | PF3D7_0304600 | sporozoite surface, dominant antigen | yes | + | - | - | + | - | - | yes [12] |
| Repetitive interspersed families of polypeptides (Rifin; RIF) | PF3D7_1040800 | variant antigen | ? | + | - | - | + | - | - | - |
| Erythrocyte binding antigen 181 (EBA181) | PF3D7_0102500 | parasite invasion | yes | + | + | + | + | + | + | - |

| | | | | | | | | | | |
|---|---|---|---|---|---|---|---|---|---|---|
| Parasitophorous vacuole membrane protein S16 (Pfs16) | PF3D7_0406200 | sexual stage/sporozoite-specific antigen | yes | + | - | + | + | - | + | yes [13] |
| Dynactin subunit 2 (DCTN2/p50) | PF3D7_1346000 | dynein anchor protein | - | + | - | - | + | - | - | - |
| Liver stage antigen 3 (LSA3) | PF3D7_0220000 | ubiquitous antigen | - | + | + | + | + | + | + | yes [14] |
| Endoplasmin (GRP94/HSP90) | PF3D7_1222300 | ER-resident chaperone | - | + | + | + | + | + | + | - |
| Reticulocyte binding protein 2 homologue b (Rh2b) | PF3D7_1335300 | merozoite invasion | yes | - | + | + | - | + | + | yes [15] |
| hypothetical protein | PF3D7_0706100 | - | ? | + | - | + | + | - | - | - |

Blue and red denote antigens enriched in PfSPZ-CVac (CQ) and PfSPZ-CVac (AP) immune sera, respectively.

SPZ, salivary gland sporozoites; LS, liver stages; BS, asexual blood stages.
